# Genome-wide Selection Scan in an Arabian Peninsula Population Identifies a *TNKS* haplotype Linked to Metabolic Traits and Hypertension

**DOI:** 10.1101/765651

**Authors:** Muthukrishnan Eaaswarkhanth, Andre Luiz Campelo dos Santos, Omer Gokcumen, Fahd Al-Mulla, Thangavel Alphonse Thanaraj

## Abstract

Despite the extreme and varying environmental conditions prevalent in the Arabian Peninsula, it has experienced several waves of human migrations following the out-of-Africa diaspora. Eventually, the inhabitants of the peninsula region adapted to the hot and dry environment. The adaptation and natural selection that shaped the extant human populations of the Arabian Peninsula region have been scarcely studied. In an attempt to explore natural selection in the region, we analyzed 662,750 variants in 583 Kuwaiti individuals. We searched for regions in the genome that display signatures of positive selection in the Kuwaiti population using an integrative approach in a conservative manner. We highlight a haplotype overlapping *TNKS* that showed strong signals of positive selection based on the results of the multiple selection tests conducted (integrated Haplotype Score, Cross Population Extended Haplotype Homozygosity, Population Branch Statistics, and log-likelihood ratio scores). Notably, the *TNKS* haplotype under selection potentially conferred a fitness advantage to the Kuwaiti ancestors for surviving in the harsh environment while posing a major health risk to present-day Kuwaitis.

## Introduction

Archaeological evidence suggests that the Arabian Peninsula played a key role during the dispersal of modern humans out-of-Africa (Cabrera et al. 2010; Rose and Petraglia 2010; Petraglia et al. 2019). Anatomically modern humans have inhabited the Arabian Peninsula since immediately after the out-of-Africa migration; therefore, the resident populations have a long and complex evolutionary history (Petraglia and Alsharekh 2003). Being at the crossroads between Africa and Eurasia, the Arabian Peninsula served as a point of interaction of human populations and trade across the region (Groucutt and Petraglia 2012). The resettlement of peoples and traders facilitated population admixture and increased genetic diversity. Several genetic studies offer insights on how the genetic ancestry, consanguinity, and admixture have structured the genetic diversity of the Arab populations. For example, uniparental genetic examinations demonstrated maternal (Abu-Amero et al. 2007; Rowold et al. 2007; Abu-Amero et al. 2008) and paternal (Abu-Amero et al. 2009; Triki-Fendri et al. 2016) genetic affinities and admixture events. Genome-wide characterization studies (Behar et al. 2010; Hunter-Zinck et al. 2010; Alsmadi et al. 2013) have elaborated the genetic structure and diversity within the peninsula as well as across the continents. In addition, a recent whole-exome–based study revealed the heterogeneous genetic structure of the Middle Eastern populations (Scott et al. 2016).

The population of the State of Kuwait exemplifies the overall heterogeneity of Middle Eastern populations (Alsmadi et al. 2013; Alsmadi et al. 2014). Kuwait is one of the seven countries located in the Arabian Peninsula on the coastal region of the Arabian Gulf. Also placed at the head of the Persian Gulf, Kuwait is bordered by Saudi Arabia and Iraq to the south and north, respectively. The ancestors of the extant Kuwaiti population were early settlers that migrated from Saudi Arabia (Alghanim 1998; Casey 2007). Until the discovery of oil, they mostly derived their livelihoods from fishing and merchant seafaring (Lienhardt 2001). The movement of populations from settlements in neighboring regions, particularly Saudi Arabia and Persia (Alenizi et al. 2008), in addition to the consequent admixture between populations and consanguinity (Yang et al. 2014), have potentially shaped the genetic diversity of the Kuwaiti population. Our previous studies revealed that the genetic structure of the Kuwait population is heterogeneous, comprising three distinct ancestral genetic backgrounds that could be linked roughly to contemporary Saudi Arabian, Persian, and Bedouin populations (Alsmadi et al. 2013; Alsmadi et al. 2014; Thareja et al. 2015).

Paleoanthropological studies have recorded the dramatic environmental transformations and extreme climatic conditions in the Arabian Peninsula over time in addition to the subsequent human dispersal into the region (Groucutt and Petraglia 2012). The extreme and varying environmental conditions could have influenced natural selection and triggered adaptation to the hot and dry desert climates (Rose and Petraglia 2010). Additionally, the ramifications of adaptive trends reported for continental populations (e.g., lactose tolerance, skin color, resistance to blood pathogens, etc.) may have implications for the health of Arabian populations. Indeed, genome-wide selection scans have revealed positive selection for lactose tolerance, as well as skin and eye color similar to those in Europeans and malaria resistance as that in Africans (Fernandes et al. 2019). Such studies provide a robust framework for investigating the potential adaptive trends in specific populations in the Arabian Peninsula. However, such focused studies are scarce (Yang et al. 2014; Fernandes et al. 2019). For example, an earlier small-scale exploration of ancestry components suggested that genetic regions associated with olfactory pathways were under natural selection in Kuwaiti populations (Yang et al. 2014). However, the study was limited to a sample of less than 50 individuals. In the present study, we build on the findings of previous studies by increasing the sample size considerably in addition to applying multiple approaches that have become available recently to identify selection in a genome-wide manner in Kuwaiti populations (Figure 1*A*).

**Fig. 1.**
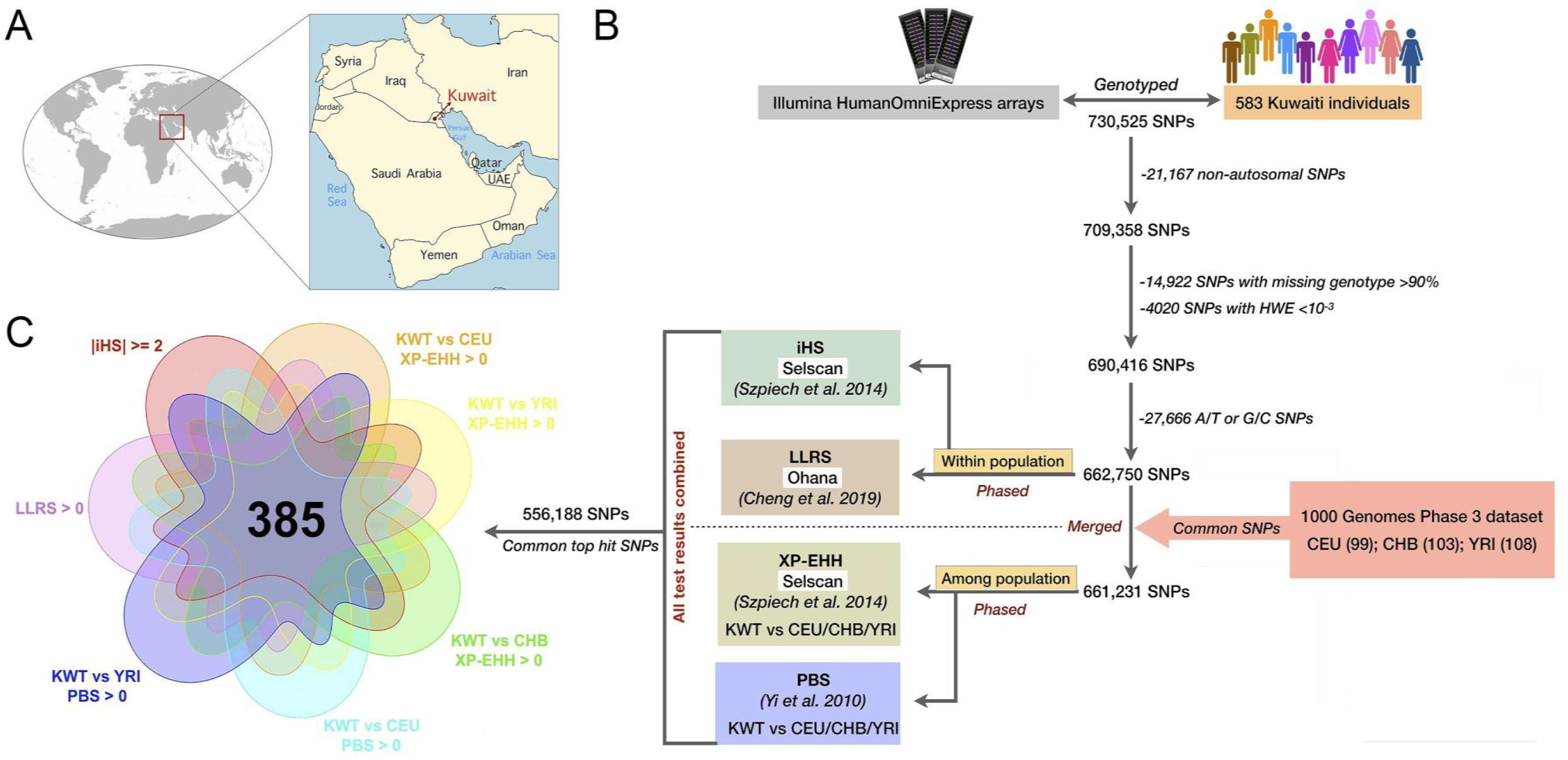
Geographic location of the genotyped individuals and materials and methods applied. **(A)** All the genotyped individuals were residents of Kuwait, in the Arabian Peninsula. **(B)** Schematic diagram of the methods, techniques, and filtering actions applied to the genome-wide genotype data. **(C)** The thresholds applied to all the selection data to identify 385 single nucleotide polymorphisms specifically under positive selection in the Kuwait population.

## Materials and Methods

### Study Samples

A total of 583 Kuwaiti individuals included in this study were selected from a larger cohort used in our previous studies (Alsmadi et al. 2013; Hebbar et al. 2017; Hebbar et al. 2018). All participants were recruited after obtaining written informed consent under protocols approved by the International Scientific Advisory Board and the Ethical Review Committee at Dasman Diabetes Institute, Kuwait. The participant recruitment, sample collection, and related procedures were conducted in accordance with the Declaration of Helsinki and detailed elsewhere (Alsmadi et al. 2013; Hebbar et al. 2017; Hebbar et al. 2018). To ensure that the study individuals are unrelated, we examined relatedness among them using PLINK (threshold PI_HAT > 0.125 *i.e.*, up to third degree relatives) and randomly removed one sample per pair of related individuals. For replication analysis we used Saudi Arabia genotype data from a published study (Fernandes et al. 2019).

### Genotyping and Quality Control

We genotyped 583 healthy unrelated Kuwaiti individuals residing in the State of Kuwait using Illumina HumanOmniExpress arrays for 730,525 single nucleotide polymorphisms (SNPs). The quality control (QC) checks and data filtering were executed in PLINK (Chang et al. 2015). The dataset was filtered through standard QC filtering to include only autosomal SNPs, specifically those that had a genotyping success rate greater than 90% and passed the Hardy-Weinberg Exact test with a p-value greater than 0.001. In addition, we eliminated strand-based ambiguous A/T and G/C SNPs. After the above filtering steps, about 662,750 SNPs remained to perform *“within population”* selection tests, integrated Haplotype Score (iHS) and log-likelihood ratio scores (LLRS) (Figure 1*B*). We merged Kuwait dataset with 1000 Genomes Project phase 3 dataset (Auton et al. 2015) yielding a combined dataset of 661,231 SNPs to conduct *“among population”* selection tests, Cross Population Extended Haplotype Homozygosity (XP-EHH) and Population Branch Statistics (PBS) (Figure 1*B*). All the four selection test results were combined, keeping only the overlapping SNPs resulting in a total of 556,188 SNPs for further examination (Figure 1*B*). To confirm whether genotyping and QC filtering criteria was adequate and that population relationships were as expected, we conducted principal component analysis on the pruned Kuwaiti population dataset combined with 1000 Genomes Project phase 3 dataset (Auton et al. 2015), using smartpca program in the EIGENSOFT package version 6.1.4 (Patterson et al. 2006; Price et al. 2006). We pruned the dataset by removing one SNP of any pair in strong pairwise linkage disequilibrium (LD) *r^2^* > 0.4 within a window of 200 SNPs (sliding the window by 25 SNPs at a time) using “indep-pairwise” option in PLINK. The SNPs were phased using Beagle (Browning and Browning 2007).

### Selection analysis

Collectively, four different selection tests that measure deviations from expected linkage disequilibrium/homozygosity (iHS, XP-EHH) and population differentiation (PBS, LLRS) distributions across the genome were performed. Each of the tests has different statistical power to detect signatures of slightly different types of selection (e.g., complete vs. incomplete sweeps) and are sensitive to the time of selection (Cadzow et al. 2014). Therefore, the integration of multiple approaches facilitated the comprehensive interrogation of the genome for selection signatures and offered a list of candidate regions that could have been evolving under non-neutral evolutionary forces.

We implemented iHS (Voight et al. 2006) and XP-EHH (Sabeti et al. 2007) tests available in selscan v1.2.0a (Szpiech and Hernandez 2014). The XP-EHH test was conducted by comparing the Kuwaiti (KWT) population with the Utah Residents with Northern and Western European Ancestry (CEU); Han Chinese from Beijing, China (CHB); and Yoruba from Ibadan, Nigeria (YRI), available from the 1000 Genomes Project phase 3 dataset (Auton et al. 2015).

We calculated PBS scores as described elsewhere (Yi et al. 2010). First, we computed SNP-wise *F*_ST_ values for all pairwise comparisons between populations – KWT, CEU, CHB, and YRI. The *F*_ST_ values were then used to calculate PBS scores in three ways: (i) between KWT and CEU using YRI as an outgroup; (ii) between KWT and CHB using YRI as an outgroup, and (iii) between KWT and YRI using CHB as an outgroup. The PBS scores were expected to pinpoint loci under selection exclusively in the Kuwaiti population.

Finally, we computed LLRS scores for positive selection using Ohana (Cheng et al. 2019). This is a recently published selection detection framework that identifies signals of positive selection through population differentiation independent of self-reported ancestry or admixture correction to group individuals into populations.

### Functional and PheWAS trait analysis

Gene functions were determined using the UCSC Genome Browser (http://genome.ucsc.edu/; last accessed May 2019) and literature searches. We performed Gene Ontology (GO) enrichment analysis using an online tool (http://geneontology.org/; last accessed May 2019) integrated with the PANTHER classification system (Mi et al. 2019). The association of the putatively selected SNPs with any phenotypic traits were detected using the PheWAS tool of the GeneATLAS database (Canela-Xandri et al. 2018). The gene expression data of the selected SNPs were obtained from the GTEx portal (https://gtexportal.org/home/; last accessed November 2019). The haplotype network was created using PopART (Leigh and Bryant 2015). The LD pattern of the positively selected seven SNPs was plotted using Haploview version 4.2 (Barrett et al. 2005).

### Code availability

All the codes related to analyses and generation of figures are available in this link.

## Results

### Principal component analysis

We performed principal component analysis (PCA) of Kuwaiti populations combined with the global populations from 1000 Genomes Project phase 3 dataset to confirm the adequacy of genotype and QC filter metrics applied and to examine population relationships. The results were consistent with the expectations. The PCA scatter plot representing the first two principal components (supplementary Figure S1) shows distinct clustering of Kuwaitis, Europeans, Africans, East Asians and South Asians revealing within and among population relationships. Consistent with our previous studies (Alsmadi et al. 2013; John et al. 2018), the placement of Kuwaiti subgroups in the proximity of their putative ancestral populations (KuwaitiB next to African cluster; KuwaitiP and KuwaitiS subgroups nearer to South Asian and European clusters) was observed.

### Integration of all selection tests scores: 385 common SNPs under selection

We interrogated 556,188 single nucleotide polymorphisms (SNPs) with all the four selection test results (Figure 1*B*, supplementary table S1). According to previously published guidelines (Cardona et al. 2014), we subsequently identified 385 SNPs, which had Cross Population Extended Haplotype Homozygosity (XP-EHH), Population Branch Statistics (PBS), and log-likelihood ratio scores (LLRS) values > 0, in addition to integrated Haplotype Score (iHS) values > 2 (Figure 1*C*, supplementary table S2). We surmised that the filtering would detect specific variants that are more likely to be true positive candidates for classical positive selection in the Kuwaiti population. As expected, the allele frequencies of the alternate alleles for the 385 SNPs were high in the Kuwaiti population in comparison with the allele frequencies for the same SNPs in the other global populations (supplementary Figure S2, supplementary table S2).

### Window-based screening and genomic regions under positive selection

We then identified the genomic locations of the 385 SNPs (Figure 2). Specifically, we speculated that if indeed selective sweeps in the Kuwaiti population explain the deviations from neutral expectations in the 385 SNPs, then they are expected to cluster within a smaller number of haplotypes. Therefore, we checked for clustering of the SNPs across the human reference genome in 100-kb windows. We identified approximately 220 windows of 100-kb in length with at least one SNP under putative selection. In addition, we observed multiple instances where adjacent 100-kb windows harbored putatively selected SNPs, which we subsequently merged into single regions (supplementary table S3).

**Fig. 2.**
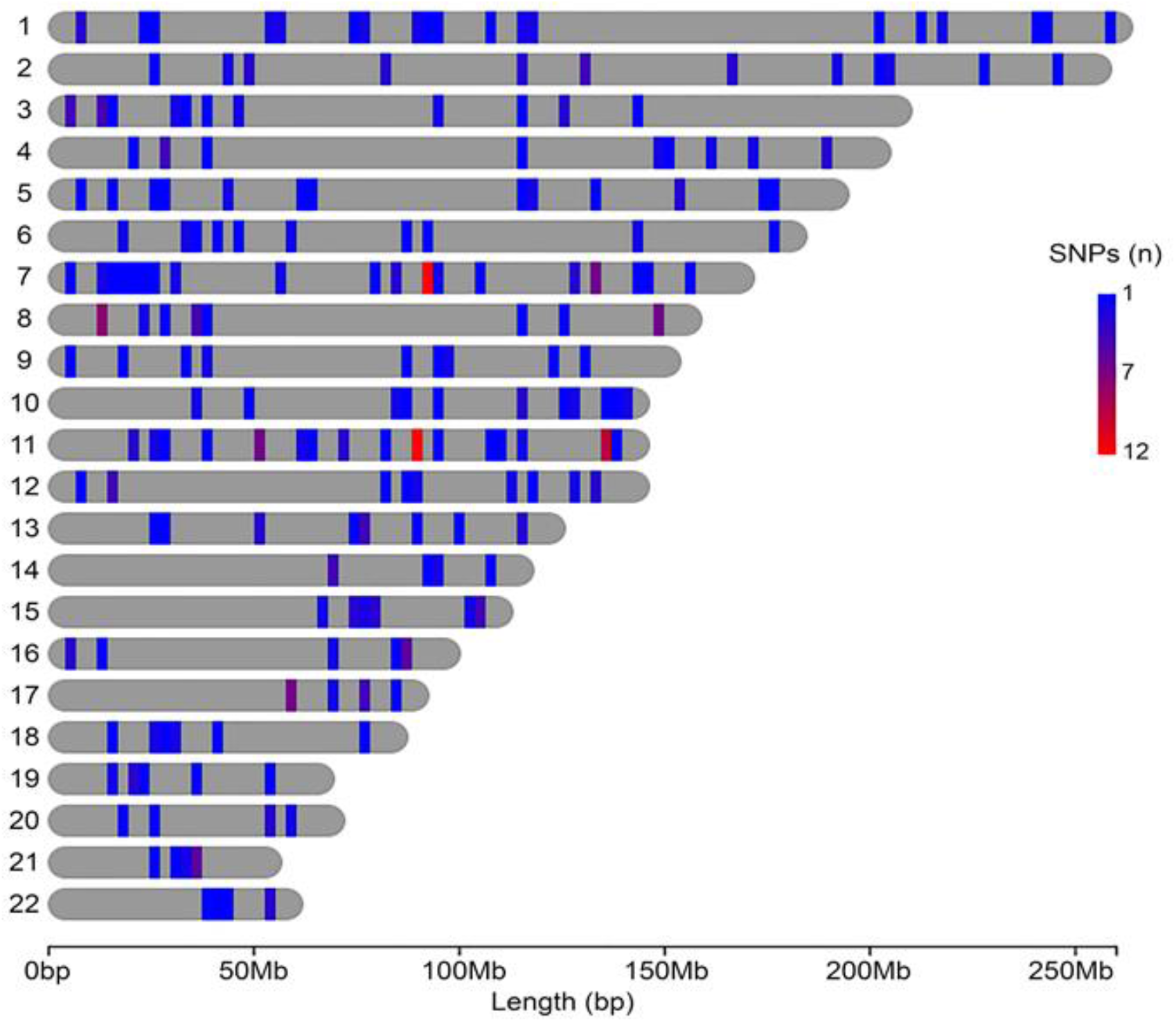
Regions in the autosomal DNA under positive selection in the Kuwait population. All the 385 SNPs under putative positive selection in the Kuwait population were assigned to 100-kb windows in the autosomal DNA. Each colored bar is a 25-Mb-segment in the chromosome representing at least one 100-kb window, and their colors reflect the number of SNPs under positive selection.

To understand the functional implications of the putatively selected regions, we identified all the genes (n = 379) inside the 220 100-kb windows and then performed a Gene Ontology enrichment analysis using all the GO annotations for *Homo sapiens* as the comparison background. After Bonferroni correction (p < 0.05), the analysis returned results associated only with a single group of biological processes: glycosaminoglycan biosynthetic process (aminoglycan biosynthetic process, aminoglycan metabolic process, glycosaminoglycan metabolic process) (supplementary table S4). There were no significant results for molecular functions and cellular components.

### TNKS haplotype related to obesity, hypertension, and asthma

To investigate further the functional impact of putatively selected haplotypes, we conducted a more thorough manual investigation of the genomic regions containing five or more SNPs identified by our selection scan. Specifically, we checked if any variant that was in linkage disequilibrium with the haplotypes had been reported in previous studies or in the UK Biobank (Sudlow et al. 2015; Bycroft et al. 2018) (supplementary table S5).

Based on the manual curation, we highlighted a ~400-kb haplotype on chromosome 8 (chr8: 9.3–9.7 Mb), which harbors seven putatively selected SNPs (Figure 3). The selection scores for all the SNPs around this haplotype region are presented in the line plots (Figure 3). To reveal the haplotype structure of the locus, we first investigated whether the seven putatively selected SNPs were in linkage disequilibrium with each other (Figure 4*B*). Indeed, a single haplotype carried the alternate alleles for all the seven SNPs (1111111) (Figure 4*A*, supplementary Figure S3). We further observed that the haplotype exhibited the highest allele frequency in the Kuwaiti population when compared with other global populations (supplementary Figure S4). In addition, within the Kuwaiti population, the subgroup of individuals with putative Saudi ancestry (Alsmadi et al. 2013) seemed to be driving up the frequency of the haplotype 1111111 (supplementary Figure S4).

**Fig. 3.**
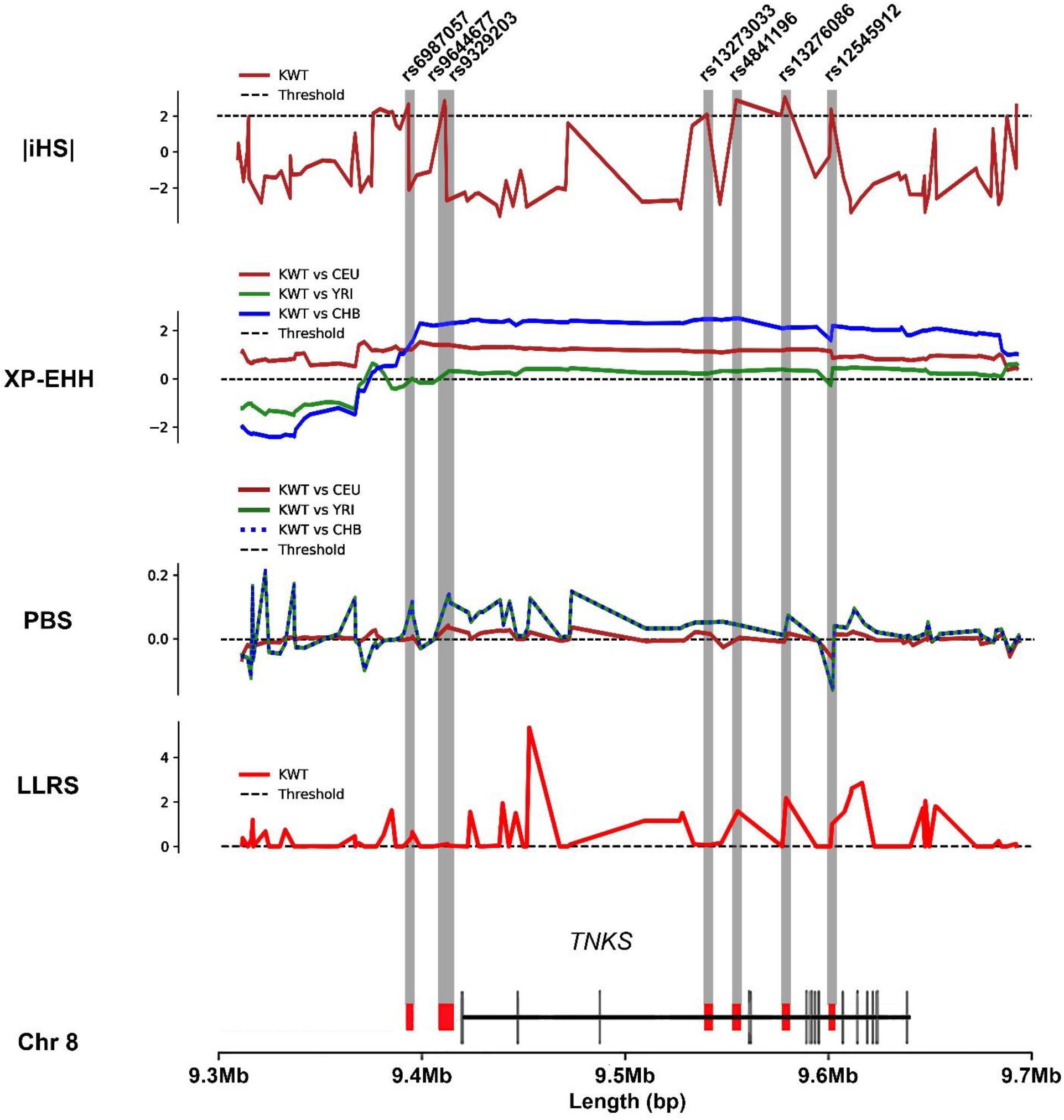
Top regions under positive selection for the Kuwait population: chr8:9.3–9.7 Mb. The values of all the selection scores for all the SNPs around the region are presented in the line plots. All the selection scores for the seven SNPs under positive selection in the region are indicated by gray bars, while their positions relative to *TNKS* are indicated by red small bars in the model at the bottom. The LD pattern for these seven SNPs is presented in fig. 4*B*.

**Fig. 4.**
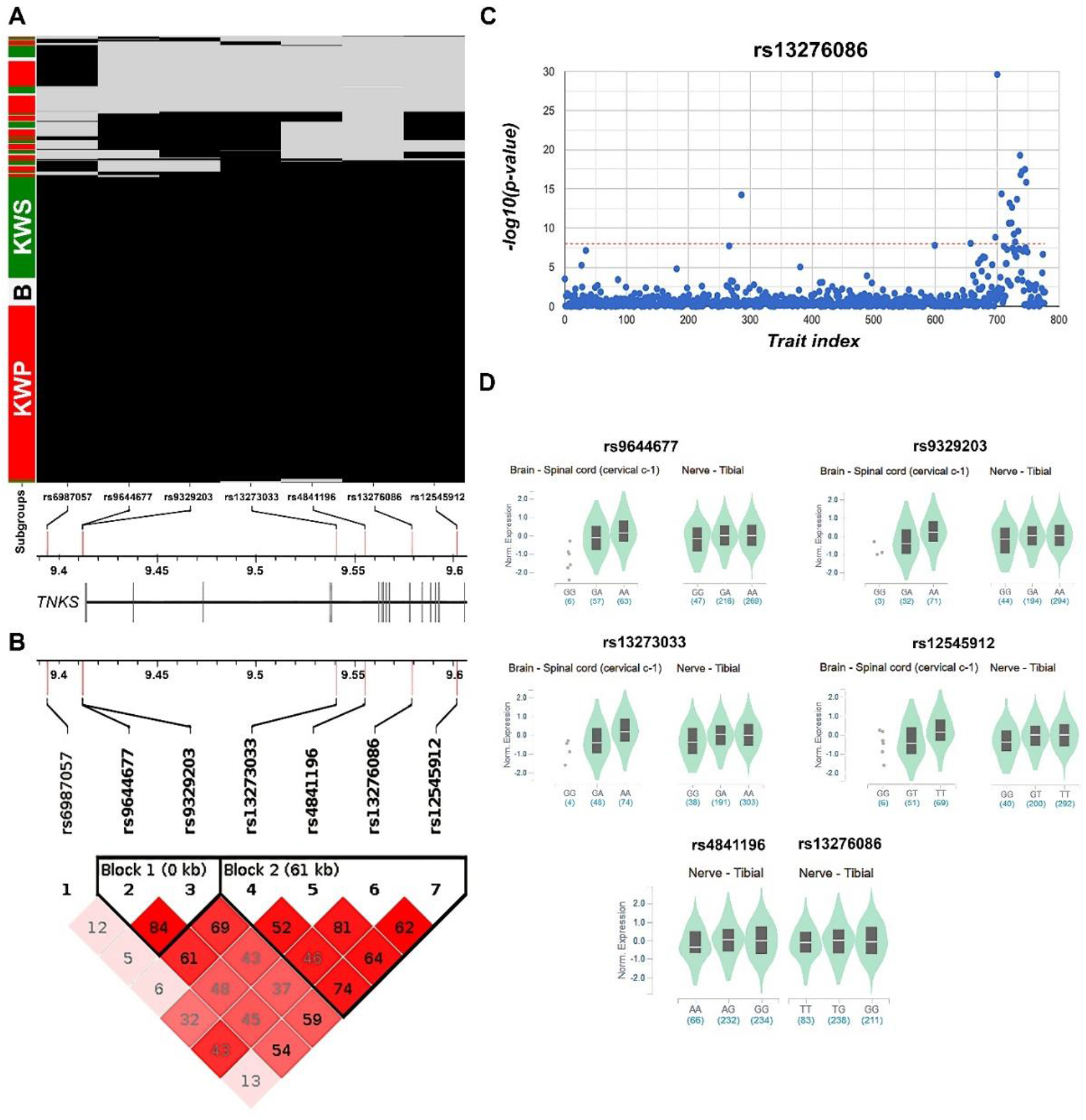
Phenotypic traits, gene expression, and haplotypes around *TNKS*. **(A)** Haplotype clustermap for the Kuwait population in the chr8:9.3–9.7-Mb region. The haplotype composed of the alternate alleles (black) of the seven SNPs under positive selection in the region is the most common even within putative subgroups, KWS - Kuwaitis with putative Saudi ancestry (green); KWP - Kuwaiti with putative Persian ancestry (red); B - Kuwaitis with putative Bedouin ancestry (gray). Each row represents one of the 583 genotyped individuals. **(B)** Linkage disequilibrium (LD) pattern of the positively selected seven SNPs. The upper panel shows the location of seven SNPs in *TNKS* and the lower panel presents the output of Haploview. Numbers in each square present the *r*^2^ value of a pairwise LD relationship between any two SNPs. **(C)** GeneATLAS PheWAS results for SNP rs13276086. In total, 17 phenotypic traits revealed an association above the threshold (-log10[p-value] > 8, red dashed line). **(D)** Violin plots of *TNKS* allele-specific eQTLs according to individual SNP genotypes in the spinal cord and the tibial nerve, in addition to the transformed fibroblasts, from the Genotype-Tissue Expression (GTEx) release V8 database. The teal region indicates the density distribution of the samples in each genotype. The white line in the box plot (black) shows the median value of the expression of each genotype.

Subsequently, we investigated the potential functional impact of the haplotype. The overall 400-kb haplotype block encompasses a single gene, the poly-ADP-ribosyltransferase 1 (*TNKS*). Three out of the seven SNPs that revealed selection signatures were located upstream of the *TNKS* while the other four were within the introns of the gene (Figure 3, 4*A*). The variation in the gene has been associated with adiposity (Lindgren et al. 2009), type 2 diabetes (Xue et al. 2018) and asthma (Ober et al. 2000). Indeed, when we searched the UK Biobank database, we found that the specific SNPs within the putatively selected haplotype that we identified using our approach were associated with metabolic disorders; for example, body mass index and limb fat mass were associated with adiposity, and eosinophil percentage was associated with asthma (George 2005) (Figure 4*C*, Table 1, supplementary table S6). Therefore, our reanalysis of the UK Biobank dataset, combined with the data obtained in both studies published previously, indicates that the specific *TNKS* haplotype is potentially associated with both metabolic traits and asthma.

**Table 1.** Summary table of the seven SNPs within the putatively selected haplotype and PheWAS traits.

| CHR | POS<sup>a</sup> | SNP | REF | ALT | Alternate Allele Frequency |  |  |  |  |  |  |  | PheWAS Trait<sup>b</sup> |
| --- | --- | --- | --- | --- | --- | --- | --- | --- | --- | --- | --- | --- | --- |
|  |  |  |  |  | KWT | SAU | AFR | AMR | EAS | EUR | SAS | ANH<sup>c</sup> |  |
| 8 | 9394053 | rs6987057 | T | G | 0.83 | 0.91 | 0.69 | 0.65 | 0.60 | 0.84 | 0.73 | — | Platelet distribution width, Body mass index (BMI), Red blood cell (erythrocyte) distribution width |
| 8 | 9411808 | rs9644677 | G | A | 0.79 | 0.87 | 0.57 | 0.79 | 0.58 | 0.75 | 0.66 | 0.62 | Hypertension |
| 8 | 9412153 | rs9329203 | G | A | 0.81 | 0.90 | 0.63 | 0.80 | 0.57 | 0.77 | 0.67 | — | Hypertension, Platelet distribution width, Standing height |
| 8 | 9540693 | rs13273033 | G | A | 0.83 | 0.92 | 0.72 | 0.82 | 0.67 | 0.78 | 0.70 | 0.69 | Hypertension, Platelet distribution width, Standing height |
| 8 | 9555092 | rs4841196 | A | G | 0.74 | 0.83 | 0.61 | 0.57 | 0.38 | 0.72 | 0.57 | — | Hypertension, Platelet distribution width, Body mass index (BMI), Red blood cell (erythrocyte) distribution width, Arm fat percentage (left), Arm fat percentage (right), Impedance of leg (left), Impedance of leg (right) |
| 8 | 9578982 | rs13276086 | T | G | 0.72 | 0.83 | 0.55 | 0.54 | 0.46 | 0.67 | 0.57 | — | Hypertension, Platelet distribution width, Standing height, Body mass index (BMI), Red blood cell (erythrocyte) distribution width, Arm fat percentage (left), Arm fat percentage (right), Impedance of leg (left), Impedance of leg (right), Arm fat mass (left), Arm fat mass (right), Cheese intake, Eosinophill percentage, Impedance of arm (right), Impedance of whole body, Leg fat mass (left), Leg fat mass (right) |
| 8 | 9601699 | rs12545912 | G | T | 0.80 | 0.90 | 0.65 | 0.84 | 0.72 | 0.77 | 0.69 | 0.62 | Hypertension, Platelet distribution width, Standing height |

KWT-Kuwaiti; SAU-Saudi Arabian; AFR-African; AMR-Admixed American; EAS-East Asian; EUR-European; SAS-South Asian; ANH-Ancient Humans.
<sup>a</sup>The SNPs positions are based on the human reference genome build GRCh37/hg19.
<sup>b</sup>Threshold: $-\log_{10}(\text{p-value}) > 8$ , PheWAS analysis provided through GeneATLAS; Individual p-values are presented in Table S6.
<sup>c</sup>ANH data from https://reich.hms.harvard.edu/downloadable-genotypes-present-day-and-ancient-dna-data-compiled-published-papers.

In addition to verifying previous associations, we observed that six out of seven putatively selected SNPs in the region were associated significantly with hypertension, as inferred from the UK Biobank’s GeneATLAS database (Canela-Xandri et al. 2018). To the best of our knowledge, this is the first report linking the region to hypertension, and considering hypertension is associated with high levels of adiposity (obesity) (Beevers et al. 2001), it further implicates the variation in *TNKS* in the pleiotropic effects on metabolism at the organismal level. In addition, expression data of *TNKS* revealed that depending on which of the seven SNPs of the haplotype a given individual carries, it is upregulated mainly in the spinal cord and tibial nerve, besides the transformed fibroblasts (Figure 4*D*). According to existing data based on expression quantitative trait loci, *TNKS* is expressed globally and mainly in brain-related tissues. The spinal cord along with the brain structures and peripheral nervous system keep up the sympathetic nervous system activity, which is involved in metabolic disorders associated with hypertension (Tanaka and Itoh 2019). Accordingly, the upregulation of *TNKS* in spinal cord or tibial nerve is crucial in contributing to the metabolic processes.

It is critical to note here that most association studies that link genetic to phenotypic variation have been conducted in people of European descent; therefore, they have a limited capacity to capture population–specific associations that are currently widespread comprehensively. However, considering the derived haplotype is extremely common in multiple populations, a more general impact could be deduced from already available databases.

### Selection in Saudi Arabia: overlap between positively selected regions linked to metabolic traits

In an effort to detect signals of positive selection in another population from the Arabian Peninsula applying our integrated conservative approach, we screened the same set of variants (662,750) genotyped from 96 Saudi Arabian individuals available from a previous study (Fernandes et al. 2019). We observed differential pattern of selection signals in Saudi Arabians (supplementary table S7). We found that different regions of the genome showed signatures of selection between Kuwaiti and Saudi Arabian populations. However, of the top 14 genomic regions under selection in Saudi Arabians, four with few common SNPs overlap with Kuwaitis. It is interesting to note that the panel of gene regions that are positively selected in Saudi Arabians also have been associated with metabolic traits (Kathiresan et al. 2007; Paterson et al. 2010) and asthma (Ober et al. 2000). For example, adiponectin levels that are correlated with obesity risk (Qi et al. 2011) and type 2 diabetes (Meigs et al. 2007). This replication analysis indicates that the hypertension related observation is specific to Kuwaiti population.

## Discussion

We identified gene regions that presented strong signals of positive selection in Kuwaiti populations using an integrative approach. Our decision to focus only on putatively selected SNPs that were identified based on multiple tests of selection made our approach highly conservative and, therefore, not prone to a high false-positive rate. For example, some of the top candidates identified using PBS (not exhibiting adequately high signals in other selection tests) cluster into a haplotype block that included *LCT*. The result was not surprising since lactose tolerance is one of the most extensively studied adaptive traits and it has been published elsewhere that *LCT* haplotypes were positively selected in the Middle Eastern populations (Enattah et al. 2008; Bayoumi et al. 2016; Liebert et al. 2017).

With our approach, we identified regions that have similar functional relevance (*e.g.*, metabolism) in Kuwaiti and Saudi Arabian populations. However, these regions do not overlap between these two populations indicating that the putative selective pressures in the region may have different ramifications in the genomes of these populations. The absence of overlapping selection signals can be attributed to the different outcomes of selection tests and an heterogeneous population structure. The extant Arabian Peninsula populations exemplify interregional genetic heterogeneity, which is evident in Saudi Arabian and Kuwaiti populations (Hunter-Zinck et al. 2010; Alsmadi et al. 2013; Tadmouri et al. 2014; Scott et al. 2016; Hajjej et al. 2018; Khubrani et al. 2018). Thus, it may not be surprising that not all four selection tests return similar results in Kuwaiti and Saudi Arabian populations. For example, using the same conservative thresholds for multiple selection tests, we were not able to replicate the selection signal for the *TNKS* haplotype in the Saudi Arabian population. Specifically, out of the seven SNPs for which we documented above-threshold signals for four different selection tests in the Kuwaiti population, four had lower iHS values than our threshold, and none of these SNPs showed increased LLRS values in the Saudi Arabian population. This result does not mean that the haplotype frequency and haplotype structure of this locus can be explained by neutrality alone in the Saudi Arabian population.

On the contrary, all seven SNPs that we highlighted in the Kuwaiti population also showed above-threshold PBS and XP-EHH values in the Saudi Arabian population (supplementary table S8). Moreover, the putatively selected *TNKS* haplotype (with all SNPs have the alternative allele) has a higher allele frequency in the Saudi Arabian population (78%) than in the Kuwaiti population (68%). In fact, it is the frequency of the Kuwaiti population sub-group with Saudi Arabian ancestry (76%) that has raised the overall occurrence of this all SNPs derived allele haplotype (1111111) in Kuwaitis (supplementary Figure S4). Thus, there is reason to believe that this haplotype is also selected in the Saudi Arabian population. The iHS test may have reduced power because the allele is at or close to fixation (Voight et al. 2006), while the LLRS may have reduced power due to a different population structure in Saudi Arabia (Cheng et al. 2019). Therefore, the lack of overlap between different populations should be considered as an outcome of multiple selection tests with different sensitivities. Overall, our study establishes a robust framework for the generation of additional adaptive hypotheses for Arabian Peninsula populations.

In the present study, we chose to highlight one 400-kb haplotype that was detected to be under positive selection based on multiple tests. The haplotype encompasses *TNKS*, and we verified its association with metabolic traits (Table 1, supplementary table S6). Indeed, we observed a considerable prevalence of the seven *TNKS* haplotype SNPs in modern continental populations and three SNPs in ancient humans (Table 1). In addition, we found that the haplotype is associated with hypertension and an increase in *TNKS* expression in the nervous system. An earlier study identified a *TNKS* intronic variant, rs6994574_G>A with signals of recent positive selection in East Asian (CHB+JPT) and African (YRI) populations from HapMap Phase 2 database based on long-range haplotype (LRH) and iHS tests (Supplementary Table 9 in International HapMap Consortium et al. 2007). This SNP information is not available in the Kuwait and Saudi Arabia datasets, as it was not included in the Illumina HumanOmniExpress arrays used for genotyping in the current study. However, rs6994574 is in strong LD with one of the seven positively selected SNPs rs13276086 in East Asians (*r*^2^ = 1) and Africans (*r*^2^ = 0.8). It is noteworthy that rs13276086 has been associated with 17 phenotypic traits according to GeneATLAS PheWAS database (Figure 4*C*, supplementary table S6). Interestingly, the alternate alleles of both the SNPs, rs6994574_A and rs13276086_G have been significantly related to hypertension with an odds ratio of 0.97 (http://geneatlas.roslin.ed.ac.uk/phewas/; Canela-Xandri et al. 2018). Another study examining telomere biology genes showed evidence of balancing selection in the *TNKS* region, consisting of 52 SNPs, by evaluating population differentiation (*F*_ST_), genetic diversity, allele frequency, and LD in global populations from HGDP-CEPH, HapMap Phases 2, and 3 databases (Mirabello et al. 2012). Notably, those 52 *TNKS* SNPs include three of the seven SNPs under putative selection that we report in the present study (rs9644677, rs13273033, rs12545912). Yet another Tajima's D statistic based investigation in individuals of Pacific Rim ancestry showed a balancing selection of 48 SNPs in the *TNKS* region (Savage et al., 2005).

The harsh desert climate of the Kuwait region could have driven the selection (Weder 2007; Young 2007). For example, a recent study speculated that natural selection for insulin resistance and the associated hypertension, in addition to increased activity in the sympathetic nervous system could have been beneficial in hunter-gatherer populations conferring a hemodynamic advantage (Lewis et al. 2019). The *TNKS* is indeed very likely to be pleiotropic. It is expressed widely across multiple organs, with especially abundant expression in the adult nervous system (Fagerberg et al. 2014). It is a poly-ADP-ribosyltransferase enzyme and its molecular function and protein partners in the cell are well understood (Kim 2018). Its activity is linked to major cellular processes including Wnt signaling pathway (Huang et al. 2009) and vesicle trafficking (Chi and Lodish 2000), as well as telomere length (Cook et al. 2002). However, little is known about the impact of haplotypic variation to *TNKS* function, and consequently the cellular and organismal phenotypes. There are multiple previous association study results that connect genetic variation within this locus with various phenotypes at a genome-wide significant p-values of <1.0E-08 (Table 1, supplementary table S6). In addition, the GWAS Catalog (https://www.ebi.ac.uk/gwas/genes/TNKS) lists several *TNKS* variants associated with multiple disorders and traits, such as multiple myeloma, neuroticism, bipolar disorder, schizophrenia, blood pressure, obesity and bone mineral density. Overall, it is plausible that the putatively adaptive haplotype has medically negative consequences that may fit a "thrifty" (Ayub et al. 2014) or "drifty" (Speakman 2008) gene scenario. However, additional work is needed to properly test this hypothesis.

The pleiotropic effects of the *TNKS* haplotypes are consistent with the speculative phenotypes described in the present study. Notably, other gene regions under potential selection have also been associated with obesity and hypertension (supplementary text, supplementary table S5). Therefore, it is plausible that the *TNKS* haplotype exemplifies a general trend in which a more rapid metabolism rate and hypertension have been selected in the Kuwaiti population, which increased the allele frequency of multiple haplotypes and conferred some degree of fitness advantage to ancestors of present-day Kuwaiti populations in the extremely dry and hot ecological environments.

In modern Kuwait, however, the effect of the *TNKS* haplotype is potentially detrimental. Indeed, hypertension and obesity are prevalent in the Kuwaiti population, affecting a staggering 25.3% and 48.2% of the population, respectively (Al-Sejari 2018). The World Health Organization has estimated that the mortality rate in Kuwait due to noncommunicable diseases is approximately 72% (https://www.who.int/nmh/publications/ncd-profiles-2018/en/; last accessed June 2019), which is alarmingly high. Such mortality and morbidity levels could be attributed largely to the drastic changes in the lifestyles and behaviors associated with westernization following oil discovery. Nevertheless, our results suggest that past adaptive trends have further predisposed Kuwaiti populations to the illnesses above at the genetic level. Overall, the mechanisms through which the *TNKS* haplotype conferred a fitness advantage and how the same haplotype predisposes the population to metabolic diseases remain fascinating areas that could be explored in future research.

## Acknowledgments

The work was supported by the Kuwait Foundation for the Advancement of Sciences research grant for Dasman Diabetes Institute (RA 2015-022). We thank the members of the National Dasman Diabetes BioBank Core Facility for sample processing and DNA extraction. We highly acknowledge Drs. Luisa Pereira and Veronica Fernandes for providing data for replication analysis. Editorial Support was provided by Dr. Diana Marouco. We thank Prashantha Hebbar for processing the raw genotype data.

## Author Contributions

T.A.T., O.G., and M.E. designed the study; M.E., and A.L.C.S., conducted most analyses; O.G., contributed significantly to the interpretation of the results; M.E., A.L.C.S., and O.G. wrote the main paper; T.A.T., and F.A-M. contributed to the writing of the paper; F.A-M. provided required resources, critically reviewed, and approved the paper; All authors reviewed the paper.

