## Supplementary Figures for "Genome-wide Selection Scan in an Arabian Peninsula Population Identifies a *TNKS* haplotype Linked to Metabolic Traits and Hypertension"

### Supplementary Tables

**Supplementary table S1.** The selection test scores and the allele frequencies of all the variants.
**Supplementary table S2.** The selection test scores and the allele frequencies of 385 variants.
**Supplementary table S3.** The 100-kb window regions and the number of SNPs under putative selection.
**Supplementary table S4.** The Gene Ontology analysis results after Bonferroni corrections.
**Supplementary table S5.** Regions under selection in the Kuwait population and associated disease phenotypes.
**Supplementary table S6.** The PheWAS traits of the seven SNPs within the putatively selected haplotype.

**Supplementary fig. S1. Density plot comparing the allele frequencies (AF) of the 385 positively selected SNPs in the Kuwait population.** In general, KWT (Kuwaiti) AFs are higher than AFR (African) and EAS (East Asian) AFs, while at some SNPs, they could be lower when compared to the EUR (European) AF and contest with the AMR (Admixed American) and SAS (South Asian) AFs for the second or third spots when they are not the highest.

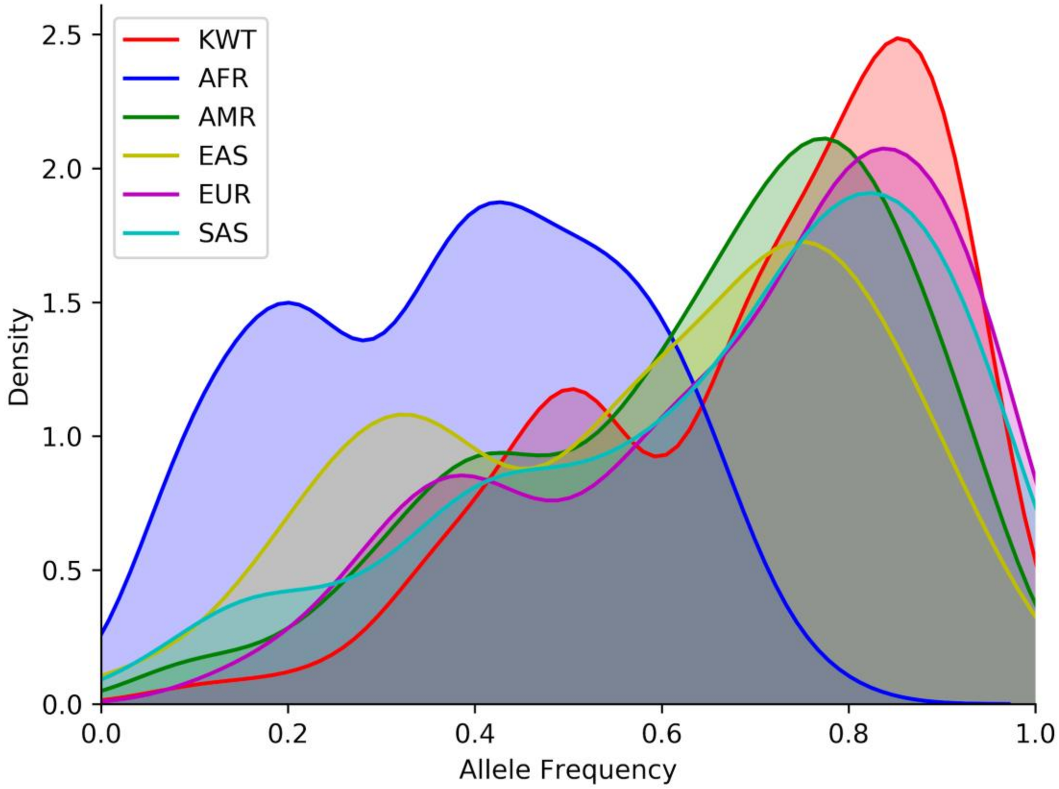

**Supplementary fig. S2. Haplotype network for the seven SNPs under positive selection for the Kuwait population in the chr8:9.3–9.7-Mb region.** To construct the network, four populations were considered, including Kuwaiti (KWT), Utah Residents with Northern and Western European Ancestry (CEU), Yoruba (YRI), and Han Chinese (CHB). Each slice in the pie charts represents the normalized number of individuals in their respective populations carrying the haplotype, while charts sizes represents the total number of individuals carrying the haplotype, limited to a maximum size for better visualization of the smallest ones. 0 = Reference Allele; 1 = Alternate Allele; 1111111 = individuals that carry the alternate alleles for all the seven SNPs.

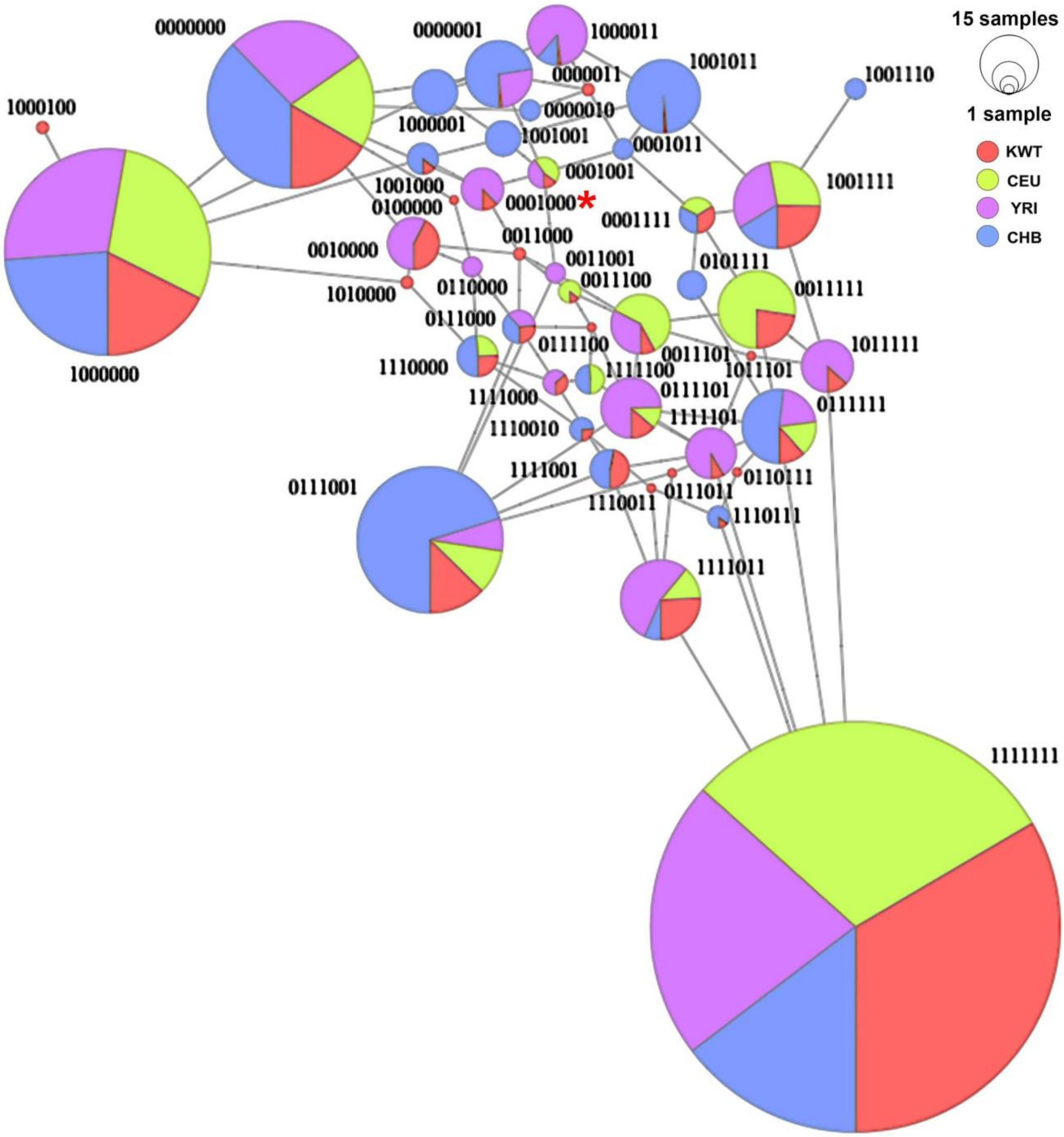

**Supplementary fig. S3. Percentages of the top five most frequent haplotypes for the chr8:9.3–9.7-Mb region SNPs among all analyzed populations and subpopulations.** (left) The KWT (Kuwaiti) population carries the haplotype 1111111 at the highest frequency, (right) while within the population, the Kuwaitis with putative Saudi ancestry (KWS) drive this haplotype frequency to higher values. KWP - Kuwaitis with putative Persian ancestry; KWB - Kuwaitis with putative Bedouin ancestry.

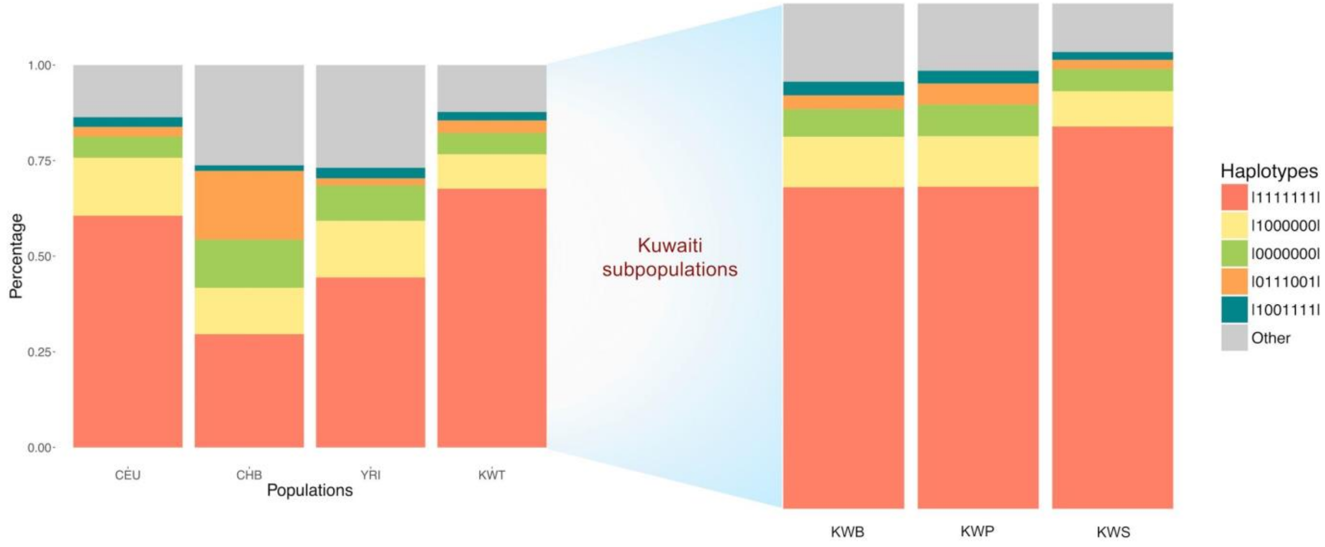

### Supplementary Text

#### Other regions under selection

Other two out of the top eight regions under positive selection for the Kuwait population, regions chr11:128.8–129 Mb and chr16:81–81.2 Mb, also revealed both diseases and traits in addition to other phenotypic associations. The first of the mentioned regions harbored six SNPs, one was intergenic while the other five affect two genes, *TP53AIP1* (1 SNP) and *ARHGAP32* (4). *ARHGAP32* was particularly interesting due to its previously reported association with low-density lipoprotein cholesterol (LDL-C) according to the UCSC Genome Browser (Kathiresan et al. 2007). The finding could also be linked to the results presented in the previous section since high-levels of the LDL-C are often associated with obesity (high BMI) and hypertension (Egan et al. 2013; Klop et al. 2013). A single phenotypic trait, mean platelet (thrombocyte) volume, is associated with the six SNPs (**supplementary table S5**). Recent studies have shown that oxidized LDL-C activates platelets (Boullier et al. 2001; Podrez et al. 2007; Magwenzi et al. 2015; Wang and Tall 2016). Therefore, we could hypothesize that the mean platelet volume in a given organism could be directly proportional to their levels of cholesterol, particularly LDL-C.

The second of the two regions, chr16:81–81.2 Mb, harbored five SNPs specifically under positive selection for the Kuwait population, affecting three genes: *ATMIN* (three SNPs), *C16orf46* (1), *GCSH* (1). Among the three genes, *C16orf46* has been associated previously with several factors, namely, attention deficit hyperactivity disorder, conduct disorder (Anney et al. 2008), body weight and measurements (Fox et al. 2007), and smoking cessation (Uhl et al. 2010). Although the first disorder seems to have little to no relation to the results presented here, the other two are consistent with our other findings. An individual’s body weight and measurements can be directly correlated with adiposity/obesity/BMI. In addition, although in recent years the discussion about the relationship between smoking and blood pressure has not been univocal (Halperin et al. 2008; Li et al. 2010), the habit is considered to play a key role in the pathophysiology of hypertension. For example, a more conclusive recent study reported that blood pressure was lower in current smokers compared to nonsmokers and former smokers, and that smoking cessation was significantly associated with an increased risk of hypertension (Li et al. 2017). Notably, Kuwait is one of the countries with increasing smoking prevalence (Reitsma et al. 2017). Smoking is a very common habit in the Kuwaiti population and most of the smokers begin the habit as early as 20 years of age (Memon et al. 2000). This can be attributed to regular social gathering activities, which are common in the Arab culture (Al-Sejari 2018).

Regarding the phenotypic associations with the SNPs in the second region, almost all the three traits, including mean reticulocyte volume, mean corpuscular volume, and mean sphered cell volume, were associated with the five SNPs, (**supplementary table S5**). The exception was the SNP harbored by *GCSH*, which was associated only with the first two of the above-mentioned traits. Recent studies have reported that hypertension (Shimizu et al. 2019)/pulse pressure (Novelli et al. 2014) in a given individual is consistently and positively correlated with reticulocyte proportion/levels. Since all the three traits are associated with red blood cells and their characteristics, we conclude that there is also an association between the phenotypes in Kuwait and our previously reported findings.
