## Supplementary Table S2-S6 for "Genome-wide Selection Scan in an Arabian Peninsula Population Identifies a *TNKS* haplotype Linked to Metabolic Traits and Hypertension"

Table S2. The selection tests scores and the allele frequency of 385 variants.

| CHR | POS | SNP | iHS | XP-EHH | | | PBS | | | LLRS | Alternate Allele Frequency | | | | | |
|---|---|---|---|---|---|---|---|---|---|---|---|---|---|---|---|---|
| | | | | KWT vs CEU | KWT vs YRI | KWT vs CHB | KWT vs CEU | KWT vs YRI | KWT vs CHB | | KWT | AFR | AMR | EAS | EUR | SAS |
| 1 | 3362256 | rs867814 | 2.688 | 0.736 | 1.550 | 1.376 | 0.017 | 0.149 | 0.149 | 0.315 | 0.82 | 0.52 | 0.75 | 0.59 | 0.84 | 0.83 |
| 1 | 3389727 | rs2185639 | 2.680 | 1.092 | 1.351 | 1.484 | 0.008 | 0.079 | 0.079 | 0.271 | 0.84 | 0.39 | 0.76 | 0.67 | 0.86 | 0.87 |
| 1 | 3393037 | rs12124163 | 2.646 | 1.130 | 1.047 | 1.494 | 0.006 | 0.061 | 0.061 | 0.395 | 0.84 | 0.36 | 0.76 | 0.68 | 0.86 | 0.86 |
| 1 | 18052645 | rs260514 | 2.422 | 1.362 | 1.621 | 2.805 | 0.047 | 0.413 | 0.413 | 0.435 | 0.94 | 0.63 | 0.93 | 0.68 | 0.96 | 0.86 |
| 1 | 22282619 | rs7529220 | 2.012 | 0.647 | 1.985 | 2.245 | 0.030 | 0.164 | 0.164 | 7.994 | 0.86 | 0.52 | 0.82 | 0.76 | 0.86 | 0.88 |
| 1 | 48295710 | rs7525331 | 2.496 | 1.704 | 2.335 | 2.291 | 0.062 | 0.447 | 0.447 | 0.220 | 0.93 | 0.58 | 0.71 | 0.59 | 0.96 | 0.85 |
| 1 | 48370525 | rs1556981 | 2.691 | 1.149 | 3.035 | 2.541 | 0.111 | 0.374 | 0.374 | 0.799 | 0.88 | 0.56 | 0.75 | 0.62 | 0.87 | 0.80 |
| 1 | 50666515 | rs2494876 | 2.540 | 1.160 | 1.861 | 1.036 | 0.005 | 0.224 | 0.224 | 1.904 | 0.89 | 0.34 | 0.88 | 0.65 | 0.90 | 0.88 |
| 1 | 50716267 | rs4437920 | 2.123 | 1.030 | 1.819 | 0.085 | 0.092 | 0.174 | 0.174 | 0.493 | 0.93 | 0.58 | 0.94 | 0.82 | 0.93 | 0.86 |
| 1 | 69053937 | rs2064375 | 2.363 | 3.151 | 1.548 | 0.221 | 0.051 | 0.007 | 0.007 | 1.036 | 0.73 | 0.37 | 0.65 | 0.72 | 0.64 | 0.70 |
| 1 | 71288661 | rs11209700 | 2.356 | 0.345 | 0.600 | 0.496 | 0.017 | 0.024 | 0.024 | 4.797 | 0.24 | 0.18 | 0.10 | 0.00 | 0.21 | 0.04 |
| 1 | 71325340 | rs6672081 | 2.133 | 0.277 | 0.366 | 0.733 | 0.012 | 0.064 | 0.064 | 4.967 | 0.25 | 0.07 | 0.10 | 0.13 | 0.22 | 0.13 |
| 1 | 84787312 | rs316662 | 2.311 | 0.713 | 1.549 | 1.078 | 0.012 | 0.174 | 0.174 | 0.146 | 0.93 | 0.64 | 0.80 | 0.85 | 0.98 | 0.92 |
| 1 | 84795283 | rs315552 | 2.402 | 0.654 | 1.533 | 0.835 | 0.007 | 0.062 | 0.062 | 0.052 | 0.88 | 0.57 | 0.77 | 0.85 | 0.91 | 0.85 |
| 1 | 87293905 | rs4282790 | 2.392 | 0.002 | 1.178 | 0.377 | 0.005 | 0.009 | 0.009 | 0.020 | 0.70 | 0.29 | 0.71 | 0.78 | 0.78 | 0.79 |
| 1 | 88941573 | rs1353796 | 2.085 | 1.739 | 1.882 | 3.341 | 0.123 | 0.060 | 0.060 | 0.347 | 0.87 | 0.68 | 0.64 | 0.74 | 0.73 | 0.82 |
| 1 | 101543389 | rs4409674 | 2.112 | 0.761 | 0.609 | 0.496 | 0.014 | 0.027 | 0.027 | 2.376 | 0.17 | 0.07 | 0.07 | 0.06 | 0.12 | 0.03 |
| 1 | 107743984 | rs4915025 | 2.224 | 0.807 | 2.232 | 0.455 | 0.008 | 0.020 | 0.020 | 4.174 | 0.94 | 0.63 | 0.96 | 0.99 | 1.00 | 0.98 |
| 1 | 108467386 | rs345306 | 2.035 | 0.874 | 0.162 | 1.015 | 0.005 | 0.099 | 0.099 | 0.966 | 0.50 | 0.17 | 0.48 | 0.33 | 0.49 | 0.36 |
| 1 | 111295674 | rs420033 | 2.346 | 1.605 | 0.143 | 0.716 | 0.020 | 0.129 | 0.129 | 6.119 | 0.34 | 0.03 | 0.25 | 0.13 | 0.28 | 0.21 |
| 1 | 111573680 | rs1282330 | 2.048 | 0.534 | 0.466 | 0.190 | 0.042 | 0.014 | 0.014 | 2.878 | 0.40 | 0.15 | 0.38 | 0.35 | 0.31 | 0.16 |
| 1 | 194251044 | rs302224 | 3.882 | 0.444 | 3.945 | 1.277 | 0.115 | 0.246 | 0.246 | 0.040 | 0.91 | 0.57 | 0.90 | 0.84 | 0.91 | 0.86 |
| 1 | 203403981 | rs12757606 | 2.272 | 0.734 | 1.421 | 1.919 | 0.021 | 0.106 | 0.106 | 0.325 | 0.51 | 0.21 | 0.42 | 0.20 | 0.48 | 0.41 |
| 1 | 208960876 | rs1166923 | 2.028 | 1.146 | 0.967 | 0.978 | 0.011 | 0.008 | 0.008 | 0.034 | 0.73 | 0.47 | 0.75 | 0.76 | 0.76 | 0.85 |
| 1 | 208972144 | rs1166900 | 2.004 | 1.408 | 0.582 | 1.376 | 0.008 | 0.018 | 0.018 | 0.017 | 0.69 | 0.36 | 0.71 | 0.59 | 0.69 | 0.78 |
| 1 | 231531627 | rs480902 | 2.066 | 2.111 | 2.636 | 1.464 | 0.000 | 0.049 | 0.049 | 0.237 | 0.73 | 0.39 | 0.63 | 0.53 | 0.78 | 0.60 |
| 1 | 231813134 | rs6541281 | 2.116 | 1.644 | 1.616 | 0.836 | 0.030 | 0.176 | 0.176 | 0.097 | 0.52 | 0.22 | 0.41 | 0.18 | 0.46 | 0.44 |
| 1 | 248138217 | rs4451578 | 2.234 | 0.259 | 1.925 | 2.349 | 0.055 | 0.236 | 0.236 | 1.750 | 0.92 | 0.56 | 0.89 | 0.80 | 0.95 | 0.98 |
| 2 | 21822998 | rs1991126 | 2.169 | 0.619 | 2.301 | 0.763 | 0.125 | 0.229 | 0.229 | 0.076 | 0.94 | 0.54 | 0.92 | 0.84 | 0.96 | 0.95 |
| 2 | 38067671 | rs766232 | 2.010 | 0.987 | 1.408 | 0.134 | 0.012 | 0.028 | 0.028 | 0.218 | 0.51 | 0.35 | 0.54 | 0.35 | 0.43 | 0.49 |
| 2 | 38068167 | rs1346746 | 2.038 | 0.973 | 1.356 | 0.123 | 0.016 | 0.040 | 0.040 | 0.230 | 0.50 | 0.30 | 0.55 | 0.35 | 0.44 | 0.49 |
| 2 | 44674450 | rs1067406 | 2.218 | 1.028 | 2.048 | 1.965 | 0.010 | 0.001 | 0.001 | 0.023 | 0.69 | 0.51 | 0.71 | 0.29 | 0.66 | 0.68 |
| 2 | 44757217 | rs1011797 | 2.007 | 1.394 | 2.002 | 0.884 | 0.023 | 0.001 | 0.001 | 0.022 | 0.71 | 0.44 | 0.57 | 0.29 | 0.64 | 0.73 |
| 2 | 44845258 | rs934777 | 2.092 | 1.810 | 1.300 | 0.949 | 0.005 | 0.000 | 0.000 | 0.237 | 0.67 | 0.40 | 0.67 | 0.32 | 0.64 | 0.59 |
| 2 | 76811778 | rs10203812 | 2.046 | 1.662 | 0.657 | 1.062 | 0.001 | 0.086 | 0.086 | 1.168 | 0.49 | 0.33 | 0.25 | 0.22 | 0.45 | 0.47 |
| 2 | 76844026 | rs1474177 | 2.070 | 1.375 | 1.457 | 1.369 | 0.003 | 0.018 | 0.018 | 0.476 | 0.55 | 0.31 | 0.38 | 0.35 | 0.55 | 0.52 |
| 2 | 76865585 | rs7566719 | 2.182 | 1.586 | 1.346 | 1.371 | 0.002 | 0.142 | 0.142 | 0.454 | 0.55 | 0.20 | 0.31 | 0.22 | 0.56 | 0.51 |
| 2 | 107728660 | rs4676318 | 2.461 | 0.535 | 1.984 | 1.095 | 0.028 | 0.170 | 0.170 | 0.011 | 0.91 | 0.66 | 0.75 | 0.79 | 0.93 | 0.92 |
| 2 | 107736937 | rs7589263 | 2.427 | 0.499 | 1.583 | 1.064 | 0.005 | 0.149 | 0.149 | 0.090 | 0.86 | 0.44 | 0.70 | 0.78 | 0.89 | 0.87 |
| 2 | 107741454 | rs6759688 | 2.398 | 0.517 | 1.546 | 1.073 | 0.009 | 0.157 | 0.157 | 0.031 | 0.87 | 0.44 | 0.70 | 0.78 | 0.89 | 0.87 |
| 2 | 123662993 | rs1527973 | 2.494 | 1.760 | 2.753 | 0.781 | 0.068 | 0.031 | 0.031 | 0.067 | 0.70 | 0.58 | 0.65 | 0.64 | 0.53 | 0.70 |
| 2 | 123730872 | rs297478 | 2.295 | 2.060 | 1.998 | 0.929 | 0.093 | 0.057 | 0.057 | 0.049 | 0.76 | 0.61 | 0.67 | 0.64 | 0.55 | 0.82 |
| 2 | 123749705 | rs2247120 | 2.407 | 2.658 | 1.033 | 1.340 | 0.019 | 0.015 | 0.015 | 0.296 | 0.77 | 0.74 | 0.64 | 0.60 | 0.55 | 0.81 |
| 2 | 123752100 | rs1395930 | 2.447 | 2.660 | 1.081 | 1.336 | 0.102 | 0.074 | 0.074 | 0.032 | 0.73 | 0.55 | 0.59 | 0.60 | 0.52 | 0.75 |
| 2 | 158283299 | rs155615 | 2.052 | 0.880 | 1.460 | 0.957 | 0.010 | 0.055 | 0.055 | 0.481 | 0.89 | 0.64 | 0.83 | 0.83 | 0.91 | 0.89 |
| 2 | 158667217 | rs10497191 | 2.953 | 0.416 | 1.515 | 0.448 | 0.021 | 0.069 | 0.069 | 5.086 | 0.78 | 0.08 | 0.81 | 0.90 | 0.88 | 0.90 |
| 2 | 158691926 | rs4233672 | 2.188 | 0.340 | 1.179 | 1.338 | 0.009 | 0.000 | 0.000 | 7.108 | 0.75 | 0.21 | 0.79 | 0.79 | 0.81 | 0.81 |
| 2 | 181716272 | rs12624291 | 2.491 | 1.147 | 1.931 | 1.199 | 0.051 | 0.035 | 0.035 | 1.270 | 0.82 | 0.32 | 0.52 | 0.75 | 0.72 | 0.88 |
| 2 | 181723942 | rs4128937 | 2.506 | 1.121 | 1.934 | 1.202 | 0.047 | 0.033 | 0.033 | 1.314 | 0.82 | 0.32 | 0.52 | 0.75 | 0.72 | 0.87 |
| 2 | 192636199 | rs4527158 | 2.025 | 0.169 | 0.572 | 0.570 | 0.004 | 0.016 | 0.016 | 0.029 | 0.52 | 0.32 | 0.42 | 0.47 | 0.49 | 0.61 |
| 2 | 194731362 | rs13385986 | 2.520 | 1.494 | 1.386 | 0.364 | 0.103 | 0.068 | 0.068 | 1.055 | 0.91 | 0.66 | 0.91 | 0.87 | 0.86 | 0.93 |
| 2 | 194749557 | rs13431912 | 2.206 | 1.475 | 1.239 | 0.366 | 0.101 | 0.066 | 0.066 | 1.357 | 0.91 | 0.64 | 0.91 | 0.87 | 0.86 | 0.93 |
| 2 | 218217618 | rs899283 | 2.250 | 0.691 | 1.774 | 0.943 | 0.074 | 0.149 | 0.149 | 0.686 | 0.90 | 0.48 | 0.79 | 0.80 | 0.90 | 0.76 |
| 2 | 234292684 | rs838715 | 2.227 | 2.396 | 1.027 | 0.839 | 0.040 | 0.063 | 0.063 | 0.250 | 0.43 | 0.15 | 0.37 | 0.36 | 0.37 | 0.25 |
| 3 | 1020877 | rs6781767 | 2.084 | 0.530 | 1.779 | 2.989 | 0.001 | 0.290 | 0.290 | 2.158 | 0.88 | 0.57 | 0.84 | 0.57 | 0.92 | 0.77 |

Table S2. The selection tests scores and the allele frequency of 385 variants - Continued

| CHR | POS | SNP | iHS | XP-EHH | | | PBS | | | LLRS | Alternate Allele Frequency | | | | | |
|---|---|---|---|---|---|---|---|---|---|---|---|---|---|---|---|---|
| | | | | KWT vs CEU | KWT vs YRI | KWT vs CHB | KWT vs CEU | KWT vs YRI | KWT vs CHB | | KWT | AFR | AMR | EAS | EUR | SAS |
| 3 | 1033790 | rs4401351 | 2.287 | 0.804 | 2.137 | 3.282 | 0.056 | 0.408 | 0.408 | 3.179 | 0.91 | 0.56 | 0.84 | 0.56 | 0.92 | 0.81 |
| 3 | 1034188 | rs9312040 | 2.226 | 0.820 | 2.260 | 3.465 | 0.054 | 0.392 | 0.392 | 2.891 | 0.91 | 0.58 | 0.84 | 0.57 | 0.92 | 0.82 |
| 3 | 1035503 | rs4686368 | 2.286 | 0.487 | 1.860 | 2.686 | 0.046 | 0.408 | 0.408 | 3.179 | 0.91 | 0.56 | 0.84 | 0.56 | 0.92 | 0.81 |
| 3 | 8947601 | rs547904 | 2.058 | 1.370 | 2.861 | 1.626 | 0.005 | 0.035 | 0.035 | 0.034 | 0.77 | 0.66 | 0.71 | 0.36 | 0.76 | 0.77 |
| 3 | 8955389 | rs373572 | 2.042 | 1.389 | 2.872 | 1.653 | 0.005 | 0.037 | 0.037 | 0.041 | 0.78 | 0.65 | 0.71 | 0.36 | 0.76 | 0.77 |
| 3 | 8956418 | rs574776 | 2.052 | 1.393 | 2.959 | 1.654 | 0.012 | 0.069 | 0.069 | 0.060 | 0.77 | 0.63 | 0.71 | 0.36 | 0.76 | 0.77 |
| 3 | 8975742 | rs447423 | 2.223 | 1.371 | 2.989 | 1.647 | 0.009 | 0.064 | 0.064 | 0.112 | 0.77 | 0.58 | 0.70 | 0.36 | 0.76 | 0.77 |
| 3 | 10649468 | rs12487529 | 2.676 | 0.678 | 0.975 | 1.896 | 0.008 | 0.117 | 0.117 | 0.003 | 0.46 | 0.23 | 0.33 | 0.03 | 0.57 | 0.24 |
| 3 | 25317646 | rs13090773 | 2.662 | 1.294 | 0.589 | 0.684 | 0.001 | 0.041 | 0.041 | 0.643 | 0.49 | 0.10 | 0.58 | 0.39 | 0.48 | 0.37 |
| 3 | 25318014 | rs4681015 | 2.525 | 1.537 | 0.866 | 0.844 | 0.009 | 0.036 | 0.036 | 0.119 | 0.52 | 0.23 | 0.61 | 0.38 | 0.49 | 0.35 |
| 3 | 29516639 | rs10510621 | 2.318 | 0.515 | 0.155 | 0.637 | 0.009 | 0.026 | 0.026 | 2.681 | 0.08 | 0.11 | 0.10 | 0.14 | 0.12 | 0.14 |
| 3 | 33103028 | rs6550202 | 2.249 | 0.658 | 1.740 | 1.201 | 0.008 | 0.034 | 0.034 | 2.630 | 0.90 | 0.40 | 0.82 | 0.92 | 0.89 | 0.93 |
| 3 | 41541196 | rs1495703 | 2.016 | 1.539 | 1.451 | 1.343 | 0.023 | 0.368 | 0.368 | 0.201 | 0.90 | 0.60 | 0.89 | 0.56 | 0.95 | 0.80 |
| 3 | 41559534 | rs1630957 | 2.042 | 1.838 | 1.374 | 1.330 | 0.020 | 0.330 | 0.330 | 0.082 | 0.90 | 0.63 | 0.89 | 0.56 | 0.95 | 0.81 |
| 3 | 88092679 | rs2033389 | 2.107 | 0.397 | 1.951 | 0.743 | 0.018 | 0.009 | 0.009 | 0.541 | 0.90 | 0.50 | 0.88 | 0.95 | 0.95 | 0.96 |
| 3 | 88101530 | rs1051436 | 2.694 | 0.360 | 1.964 | 0.970 | 0.016 | 0.025 | 0.025 | 0.955 | 0.82 | 0.43 | 0.84 | 0.83 | 0.84 | 0.91 |
| 3 | 109799304 | rs1358018 | 2.881 | 0.325 | 2.349 | 0.335 | 0.051 | 0.123 | 0.123 | 3.540 | 0.93 | 0.64 | 0.87 | 0.90 | 0.96 | 0.97 |
| 3 | 118865332 | rs11550908 | 2.078 | 0.425 | 0.158 | 0.530 | 0.005 | 0.012 | 0.012 | 0.086 | 0.11 | 0.13 | 0.07 | 0.24 | 0.15 | 0.11 |
| 3 | 118865609 | rs10934483 | 2.078 | 0.425 | 0.158 | 0.530 | 0.005 | 0.012 | 0.012 | 0.086 | 0.11 | 0.13 | 0.07 | 0.24 | 0.15 | 0.11 |
| 3 | 118916627 | rs12632242 | 2.010 | 0.306 | 0.648 | 1.417 | 0.040 | 0.091 | 0.091 | 1.603 | 0.40 | 0.08 | 0.31 | 0.22 | 0.30 | 0.17 |
| 3 | 137544378 | rs149999 | 2.113 | 0.375 | 0.839 | 0.030 | 0.077 | 0.119 | 0.119 | 0.806 | 0.53 | 0.18 | 0.55 | 0.67 | 0.59 | 0.65 |
| 4 | 16599234 | rs1496747 | 2.234 | 3.908 | 1.440 | 2.810 | 0.034 | 0.008 | 0.008 | 2.740 | 0.68 | 0.51 | 0.54 | 0.35 | 0.53 | 0.48 |
| 4 | 24193409 | rs6822382 | 2.396 | 0.408 | 2.646 | 1.225 | 0.004 | 0.078 | 0.078 | 3.666 | 0.83 | 0.35 | 0.82 | 0.75 | 0.84 | 0.84 |
| 4 | 24199845 | rs1397490 | 2.763 | 0.488 | 2.590 | 1.263 | 0.022 | 0.099 | 0.099 | 3.149 | 0.82 | 0.42 | 0.80 | 0.74 | 0.83 | 0.84 |
| 4 | 24205824 | rs13131294 | 2.602 | 0.496 | 2.558 | 1.248 | 0.041 | 0.042 | 0.042 | 2.029 | 0.83 | 0.51 | 0.82 | 0.83 | 0.83 | 0.84 |
| 4 | 24212477 | rs9684046 | 2.675 | 0.499 | 2.457 | 1.277 | 0.022 | 0.102 | 0.102 | 3.133 | 0.82 | 0.43 | 0.80 | 0.74 | 0.83 | 0.83 |
| 4 | 32628884 | rs755773 | 2.930 | 1.520 | 2.614 | 1.737 | 0.054 | 0.143 | 0.143 | 0.292 | 0.85 | 0.54 | 0.79 | 0.67 | 0.82 | 0.79 |
| 4 | 108681819 | rs4585383 | 2.169 | 1.026 | 2.334 | 0.282 | 0.029 | 0.082 | 0.082 | 1.094 | 0.86 | 0.45 | 0.80 | 0.78 | 0.84 | 0.91 |
| 4 | 140637815 | rs13112141 | 2.233 | 1.581 | 1.131 | 2.596 | 0.042 | 0.064 | 0.064 | 0.824 | 0.49 | 0.20 | 0.26 | 0.41 | 0.43 | 0.50 |
| 4 | 140650730 | rs2246759 | 2.125 | 1.843 | 1.096 | 2.944 | 0.027 | 0.185 | 0.185 | 1.258 | 0.51 | 0.18 | 0.28 | 0.25 | 0.45 | 0.46 |
| 4 | 142916958 | rs1995960 | 2.336 | 0.079 | 0.549 | 0.208 | 0.055 | 0.033 | 0.033 | 0.003 | 0.50 | 0.08 | 0.51 | 0.64 | 0.63 | 0.52 |
| 4 | 152855624 | rs19412 | 2.176 | 1.047 | 1.668 | 0.179 | 0.001 | 0.003 | 0.003 | 0.263 | 0.66 | 0.34 | 0.60 | 0.66 | 0.68 | 0.62 |
| 4 | 153548037 | rs3922934 | 2.149 | 0.732 | 1.415 | 0.312 | 0.049 | 0.030 | 0.030 | 0.375 | 0.92 | 0.64 | 0.95 | 0.92 | 0.94 | 0.94 |
| 4 | 162827306 | rs7666181 | 2.398 | 0.935 | 0.552 | 1.483 | 0.017 | 0.018 | 0.018 | 0.239 | 0.67 | 0.42 | 0.55 | 0.52 | 0.53 | 0.47 |
| 4 | 162904939 | rs1483549 | 2.027 | 0.553 | 0.293 | 1.795 | 0.030 | 0.048 | 0.048 | 0.392 | 0.83 | 0.54 | 0.83 | 0.77 | 0.82 | 0.83 |
| 4 | 180014056 | rs11131908 | 2.132 | 0.706 | 2.108 | 2.222 | 0.021 | 0.059 | 0.059 | 2.771 | 0.69 | 0.54 | 0.61 | 0.47 | 0.63 | 0.59 |
| 4 | 180014143 | rs1451407 | 2.275 | 0.708 | 1.935 | 2.223 | 0.007 | 0.027 | 0.027 | 4.444 | 0.68 | 0.43 | 0.60 | 0.47 | 0.63 | 0.59 |
| 4 | 180095990 | rs2383273 | 2.649 | 0.341 | 2.153 | 2.562 | 0.000 | 0.003 | 0.003 | 0.019 | 0.71 | 0.31 | 0.73 | 0.75 | 0.77 | 0.67 |
| 5 | 3333760 | rs7718013 | 2.160 | 1.012 | 1.921 | 1.443 | 0.078 | 0.105 | 0.105 | 0.048 | 0.93 | 0.67 | 0.75 | 0.88 | 0.94 | 0.88 |
| 5 | 11571940 | rs2727588 | 2.351 | 1.869 | 1.493 | 2.402 | 0.002 | 0.051 | 0.051 | 0.071 | 0.67 | 0.73 | 0.50 | 0.17 | 0.72 | 0.44 |
| 5 | 20786582 | rs10941855 | 2.394 | 1.433 | 2.158 | 0.368 | 0.004 | 0.031 | 0.031 | 5.656 | 0.89 | 0.34 | 0.80 | 0.88 | 0.93 | 0.92 |
| 5 | 24643756 | rs1395026 | 2.541 | 1.481 | 3.984 | 2.819 | 0.112 | 0.149 | 0.149 | 0.738 | 0.87 | 0.52 | 0.88 | 0.82 | 0.87 | 0.93 |
| 5 | 38103062 | rs2453341 | 2.104 | 0.549 | 1.789 | 1.911 | 0.008 | 0.153 | 0.153 | 0.040 | 0.82 | 0.50 | 0.73 | 0.70 | 0.84 | 0.84 |
| 5 | 38104956 | rs2471068 | 2.174 | 0.590 | 1.591 | 2.038 | 0.029 | 0.197 | 0.197 | 1.349 | 0.93 | 0.73 | 0.96 | 0.79 | 0.96 | 0.97 |
| 5 | 54737008 | rs11741874 | 2.022 | 1.391 | 1.891 | 1.157 | 0.001 | 0.048 | 0.048 | 0.949 | 0.69 | 0.40 | 0.57 | 0.50 | 0.66 | 0.62 |
| 5 | 57244070 | rs1378244 | 2.024 | 0.720 | 0.911 | 1.986 | 0.001 | 0.013 | 0.013 | 0.559 | 0.67 | 0.59 | 0.66 | 0.47 | 0.64 | 0.57 |
| 5 | 108922023 | rs902505 | 2.490 | 2.128 | 1.157 | 0.266 | 0.008 | 0.012 | 0.012 | 0.265 | 0.67 | 0.32 | 0.76 | 0.59 | 0.69 | 0.78 |
| 5 | 110501681 | rs244515 | 2.566 | 0.626 | 1.332 | 0.807 | 0.015 | 0.042 | 0.042 | 6.340 | 0.34 | 0.16 | 0.39 | 0.27 | 0.32 | 0.40 |
| 5 | 110544623 | rs7709420 | 2.259 | 0.443 | 0.369 | 0.367 | 0.019 | 0.015 | 0.015 | 6.054 | 0.30 | 0.18 | 0.32 | 0.27 | 0.26 | 0.28 |
| 5 | 125649162 | rs17527858 | 2.958 | 0.166 | 0.240 | 0.254 | 0.031 | 0.117 | 0.117 | 1.631 | 0.22 | 0.04 | 0.12 | 0.00 | 0.16 | 0.03 |
| 5 | 145202461 | rs6898776 | 2.418 | 0.323 | 2.659 | 0.826 | 0.019 | 0.071 | 0.071 | 1.050 | 0.57 | 0.17 | 0.57 | 0.28 | 0.69 | 0.51 |
| 5 | 145205480 | rs340046 | 2.325 | 0.335 | 2.666 | 0.918 | 0.022 | 0.049 | 0.049 | 0.843 | 0.58 | 0.21 | 0.58 | 0.31 | 0.69 | 0.51 |
| 5 | 145784922 | rs11747475 | 2.052 | 0.863 | 0.836 | 0.585 | 0.015 | 0.010 | 0.010 | 0.237 | 0.77 | 0.40 | 0.79 | 0.71 | 0.78 | 0.85 |
| 5 | 165595428 | rs11741022 | 2.017 | 0.939 | 2.610 | 2.218 | 0.013 | 0.082 | 0.082 | 1.334 | 0.86 | 0.34 | 0.70 | 0.71 | 0.88 | 0.90 |
| 5 | 165978860 | rs2964302 | 2.699 | 0.769 | 1.228 | 0.961 | 0.001 | 0.001 | 0.001 | 0.104 | 0.70 | 0.19 | 0.79 | 0.69 | 0.79 | 0.74 |
| 6 | 14864129 | rs1258993 | 2.627 | 0.179 | 2.405 | 0.109 | 0.096 | 0.020 | 0.020 | 0.924 | 0.89 | 0.46 | 0.83 | 0.93 | 0.89 | 0.89 |

Table S2. The selection tests scores and the allele frequency of 385 variants - Continued

| CHR | POS | SNP | iHS | XP-EHH | | | PBS | | | LLRS | Alternate Allele Frequency | | | | | |
|---|---|---|---|---|---|---|---|---|---|---|---|---|---|---|---|---|
| | | | | KWT vs CEU | KWT vs YRI | KWT vs CHB | KWT vs CEU | KWT vs YRI | KWT vs CHB | | KWT | AFR | AMR | EAS | EUR | SAS |
| 6 | 14884524 | rs373633 | 2.460 | 0.179 | 2.299 | 0.056 | 0.094 | 0.004 | 0.004 | 0.158 | 0.89 | 0.56 | 0.83 | 0.94 | 0.89 | 0.90 |
| 6 | 28579471 | rs9501180 | 2.690 | 0.167 | 1.175 | 1.157 | 0.006 | 0.126 | 0.126 | 0.332 | 0.49 | 0.07 | 0.44 | 0.33 | 0.56 | 0.45 |
| 6 | 28645170 | rs9393921 | 2.137 | 0.190 | 0.583 | 0.966 | 0.002 | 0.144 | 0.144 | 0.469 | 0.50 | 0.07 | 0.43 | 0.32 | 0.55 | 0.43 |
| 6 | 32286102 | rs531094 | 2.073 | 0.025 | 0.439 | 1.053 | 0.013 | 0.027 | 0.027 | 0.068 | 0.52 | 0.15 | 0.51 | 0.36 | 0.37 | 0.38 |
| 6 | 35004819 | rs12525532 | 2.571 | 0.400 | 0.340 | 0.295 | 0.009 | 0.052 | 0.052 | 0.057 | 0.40 | 0.16 | 0.38 | 0.27 | 0.39 | 0.43 |
| 6 | 40890792 | rs1929791 | 2.354 | 1.336 | 1.231 | 0.960 | 0.008 | 0.004 | 0.004 | 0.257 | 0.66 | 0.38 | 0.55 | 0.62 | 0.58 | 0.43 |
| 6 | 53478869 | rs10948754 | 2.472 | 1.317 | 1.774 | 0.963 | 0.028 | 0.052 | 0.052 | 0.576 | 0.45 | 0.25 | 0.51 | 0.30 | 0.38 | 0.18 |
| 6 | 53479069 | rs10948755 | 2.532 | 1.331 | 2.022 | 0.980 | 0.028 | 0.052 | 0.052 | 0.576 | 0.45 | 0.25 | 0.51 | 0.30 | 0.38 | 0.18 |
| 6 | 81352611 | rs2444818 | 2.389 | 1.155 | 1.605 | 0.780 | 0.004 | 0.067 | 0.067 | 0.425 | 0.44 | 0.11 | 0.46 | 0.30 | 0.38 | 0.27 |
| 6 | 84971363 | rs6910017 | 2.374 | 0.457 | 1.535 | 1.687 | 0.010 | 0.024 | 0.024 | 0.058 | 0.72 | 0.49 | 0.58 | 0.44 | 0.66 | 0.74 |
| 6 | 135739062 | rs2614290 | 2.008 | 0.716 | 2.174 | 1.624 | 0.112 | 0.087 | 0.087 | 3.573 | 0.93 | 0.50 | 0.89 | 0.97 | 0.96 | 0.94 |
| 6 | 135899344 | rs2092556 | 2.027 | 0.209 | 2.313 | 1.089 | 0.046 | 0.025 | 0.025 | 6.051 | 0.90 | 0.46 | 0.88 | 0.96 | 0.95 | 0.94 |
| 6 | 167238612 | rs1202621 | 2.004 | 0.147 | 2.565 | 2.044 | 0.030 | 0.018 | 0.018 | 0.594 | 0.71 | 0.49 | 0.55 | 0.31 | 0.65 | 0.58 |
| 7 | 1987092 | rs10237340 | 2.126 | 0.182 | 0.661 | 0.020 | 0.103 | 0.140 | 0.140 | 0.184 | 0.94 | 0.64 | 0.83 | 0.90 | 0.96 | 0.91 |
| 7 | 8249663 | rs1981715 | 2.233 | 1.197 | 1.498 | 1.338 | 0.045 | 0.130 | 0.130 | 0.018 | 0.87 | 0.64 | 0.80 | 0.67 | 0.87 | 0.90 |
| 7 | 8550648 | rs6978582 | 2.421 | 0.385 | 0.304 | 0.253 | 0.020 | 0.068 | 0.068 | 1.544 | 0.36 | 0.20 | 0.28 | 0.13 | 0.30 | 0.20 |
| 7 | 12119520 | rs17648860 | 2.515 | 1.657 | 0.394 | 0.296 | 0.015 | 0.087 | 0.087 | 1.955 | 0.27 | 0.12 | 0.12 | 0.10 | 0.24 | 0.11 |
| 7 | 14453061 | rs891808 | 2.189 | 1.294 | 1.786 | 0.912 | 0.054 | 0.106 | 0.106 | 0.015 | 0.86 | 0.63 | 0.66 | 0.66 | 0.85 | 0.71 |
| 7 | 16825580 | rs4629751 | 2.771 | 2.070 | 1.102 | 2.001 | 0.005 | 0.021 | 0.021 | 0.311 | 0.76 | 0.29 | 0.75 | 0.58 | 0.72 | 0.72 |
| 7 | 19447111 | rs4721765 | 2.388 | 0.637 | 1.769 | 2.360 | 0.002 | 0.156 | 0.156 | 4.644 | 0.84 | 0.49 | 0.69 | 0.72 | 0.88 | 0.90 |
| 7 | 21286643 | rs4548059 | 2.300 | 1.172 | 0.123 | 1.091 | 0.018 | 0.019 | 0.019 | 2.111 | 0.38 | 0.24 | 0.45 | 0.27 | 0.37 | 0.26 |
| 7 | 27182245 | rs3757640 | 3.046 | 2.537 | 1.677 | 0.542 | 0.008 | 0.015 | 0.015 | 0.491 | 0.77 | 0.36 | 0.56 | 0.60 | 0.71 | 0.75 |
| 7 | 27196113 | rs2301721 | 2.655 | 2.094 | 1.403 | 0.430 | 0.004 | 0.028 | 0.028 | 4.241 | 0.79 | 0.10 | 0.65 | 0.86 | 0.83 | 0.83 |
| 7 | 50758245 | rs933360 | 2.582 | 0.473 | 1.232 | 0.617 | 0.002 | 0.003 | 0.003 | 1.030 | 0.76 | 0.25 | 0.68 | 0.75 | 0.75 | 0.68 |
| 7 | 50791579 | rs6943153 | 2.198 | 0.497 | 0.957 | 0.429 | 0.002 | 0.030 | 0.030 | 5.314 | 0.70 | 0.25 | 0.56 | 0.76 | 0.70 | 0.67 |
| 7 | 73599917 | rs150875 | 2.179 | 0.820 | 1.148 | 0.267 | 0.006 | 0.065 | 0.065 | 0.120 | 0.93 | 0.66 | 0.92 | 0.93 | 0.97 | 0.92 |
| 7 | 78872516 | rs1406160 | 2.048 | 2.246 | 0.559 | 1.796 | 0.014 | 0.026 | 0.026 | 1.302 | 0.56 | 0.27 | 0.40 | 0.26 | 0.61 | 0.42 |
| 7 | 79057761 | rs10485959 | 2.251 | 2.323 | 0.926 | 1.609 | 0.047 | 0.030 | 0.030 | 1.430 | 0.43 | 0.11 | 0.36 | 0.25 | 0.35 | 0.33 |
| 7 | 79085543 | rs4730857 | 2.118 | 2.292 | 1.890 | 1.585 | 0.027 | 0.054 | 0.054 | 1.828 | 0.51 | 0.27 | 0.39 | 0.29 | 0.44 | 0.34 |
| 7 | 86767689 | rs7812191 | 2.487 | 0.375 | 2.308 | 1.425 | 0.039 | 0.243 | 0.243 | 0.390 | 0.82 | 0.53 | 0.62 | 0.42 | 0.82 | 0.75 |
| 7 | 86781423 | rs2286267 | 2.650 | 0.349 | 2.373 | 1.434 | 0.058 | 0.219 | 0.219 | 0.358 | 0.81 | 0.52 | 0.66 | 0.54 | 0.79 | 0.76 |
| 7 | 86847321 | rs10245036 | 2.481 | 0.215 | 2.409 | 1.066 | 0.044 | 0.203 | 0.203 | 0.085 | 0.80 | 0.52 | 0.66 | 0.53 | 0.79 | 0.75 |
| 7 | 86995999 | rs7802658 | 2.605 | 0.828 | 2.537 | 0.865 | 0.001 | 0.008 | 0.008 | 0.050 | 0.81 | 0.37 | 0.76 | 0.76 | 0.83 | 0.71 |
| 7 | 86997694 | rs10247098 | 2.570 | 0.820 | 2.843 | 0.888 | 0.006 | 0.019 | 0.019 | 0.029 | 0.81 | 0.41 | 0.77 | 0.75 | 0.83 | 0.71 |
| 7 | 86998335 | rs3789250 | 2.570 | 0.820 | 2.832 | 0.888 | 0.007 | 0.020 | 0.020 | 0.029 | 0.81 | 0.42 | 0.77 | 0.75 | 0.83 | 0.71 |
| 7 | 87022596 | rs2072207 | 2.665 | 0.835 | 2.935 | 1.164 | 0.009 | 0.048 | 0.048 | 0.336 | 0.82 | 0.41 | 0.76 | 0.71 | 0.82 | 0.72 |
| 7 | 87030903 | rs2097937 | 2.645 | 1.002 | 3.104 | 1.206 | 0.027 | 0.070 | 0.070 | 0.433 | 0.84 | 0.42 | 0.75 | 0.71 | 0.81 | 0.72 |
| 7 | 87031809 | rs1526090 | 2.607 | 1.002 | 3.104 | 1.206 | 0.029 | 0.070 | 0.070 | 0.368 | 0.84 | 0.42 | 0.75 | 0.71 | 0.81 | 0.72 |
| 7 | 87033948 | rs12154399 | 2.719 | 0.992 | 3.102 | 1.200 | 0.026 | 0.069 | 0.069 | 0.712 | 0.83 | 0.42 | 0.75 | 0.71 | 0.81 | 0.71 |
| 7 | 87043708 | rs4148830 | 2.682 | 0.942 | 3.064 | 1.169 | 0.028 | 0.069 | 0.069 | 0.410 | 0.84 | 0.42 | 0.75 | 0.71 | 0.81 | 0.72 |
| 7 | 87053150 | rs31667 | 2.210 | 0.282 | 2.726 | 1.422 | 0.010 | 0.158 | 0.158 | 0.389 | 0.85 | 0.50 | 0.79 | 0.69 | 0.84 | 0.73 |
| 7 | 89373249 | rs12667937 | 2.454 | 1.808 | 1.664 | 0.492 | 0.021 | 0.025 | 0.025 | 0.809 | 0.69 | 0.48 | 0.71 | 0.59 | 0.69 | 0.44 |
| 7 | 89410098 | rs12704484 | 2.200 | 1.821 | 1.837 | 0.549 | 0.027 | 0.009 | 0.009 | 1.834 | 0.76 | 0.35 | 0.69 | 0.73 | 0.71 | 0.49 |
| 7 | 98005398 | rs2107717 | 2.072 | 0.399 | 2.425 | 2.440 | 0.053 | 0.028 | 0.028 | 0.088 | 0.52 | 0.22 | 0.60 | 0.44 | 0.68 | 0.29 |
| 7 | 119944579 | rs12706276 | 3.090 | 1.723 | 4.512 | 0.400 | 0.001 | 0.008 | 0.008 | 2.229 | 0.81 | 0.20 | 0.79 | 0.86 | 0.81 | 0.91 |
| 7 | 119955585 | rs2402526 | 3.411 | 1.876 | 4.298 | 0.712 | 0.004 | 0.034 | 0.034 | 3.690 | 0.81 | 0.15 | 0.78 | 0.86 | 0.81 | 0.91 |
| 7 | 120010469 | rs6943092 | 2.816 | 1.879 | 4.307 | 0.733 | 0.133 | 0.114 | 0.114 | 0.048 | 0.94 | 0.48 | 0.93 | 0.92 | 0.91 | 0.94 |
| 7 | 125961043 | rs485 | 2.674 | 2.939 | 1.065 | 1.564 | 0.094 | 0.049 | 0.049 | 0.232 | 0.50 | 0.14 | 0.40 | 0.39 | 0.39 | 0.44 |
| 7 | 125992885 | rs10954109 | 2.632 | 2.816 | 0.804 | 1.589 | 0.058 | 0.035 | 0.035 | 0.170 | 0.44 | 0.15 | 0.38 | 0.37 | 0.36 | 0.36 |
| 7 | 126037381 | rs12539616 | 2.968 | 2.733 | 1.889 | 1.464 | 0.029 | 0.002 | 0.002 | 0.476 | 0.47 | 0.34 | 0.42 | 0.45 | 0.37 | 0.42 |
| 7 | 126073820 | rs1419458 | 2.345 | 2.976 | 2.003 | 1.498 | 0.021 | 0.015 | 0.015 | 0.635 | 0.54 | 0.36 | 0.46 | 0.38 | 0.42 | 0.43 |
| 7 | 126079144 | rs712723 | 2.253 | 2.862 | 1.911 | 1.432 | 0.020 | 0.015 | 0.015 | 0.568 | 0.54 | 0.36 | 0.46 | 0.38 | 0.42 | 0.43 |
| 7 | 126154236 | rs2283065 | 2.864 | 2.757 | 2.349 | 2.006 | 0.023 | 0.048 | 0.048 | 0.051 | 0.44 | 0.02 | 0.26 | 0.26 | 0.36 | 0.40 |
| 7 | 135405904 | rs6948713 | 2.107 | 1.614 | 1.343 | 1.689 | 0.041 | 0.109 | 0.109 | 3.565 | 0.48 | 0.20 | 0.25 | 0.30 | 0.40 | 0.28 |
| 7 | 137669111 | rs12707374 | 2.058 | 1.098 | 0.457 | 0.615 | 0.006 | 0.009 | 0.009 | 0.830 | 0.82 | 0.29 | 0.76 | 0.76 | 0.77 | 0.90 |
| 7 | 147581989 | rs17236183 | 2.023 | 0.248 | 0.472 | 0.239 | 0.007 | 0.079 | 0.079 | 0.856 | 0.39 | 0.07 | 0.41 | 0.27 | 0.35 | 0.31 |

| CHR | POS | SNP | iHS | XP-EHH | | | PBS | | | LLRS | Alternate Allele Frequency | | | | | |
|---|---|---|---|---|---|---|---|---|---|---|---|---|---|---|---|---|
| | | | | KWT vs CEU | KWT vs YRI | KWT vs CHB | KWT vs CEU | KWT vs YRI | KWT vs CHB | | KWT | AFR | AMR | EAS | EUR | SAS |
| 8 | 9394053 | rs6987057 | 2.628 | 1.247 | 0.034 | 1.567 | 0.004 | 0.119 | 0.119 | 0.414 | 0.83 | 0.69 | 0.65 | 0.60 | 0.84 | 0.73 |
| 8 | 9411808 | rs9644677 | 2.820 | 1.437 | 0.303 | 2.312 | 0.044 | 0.139 | 0.139 | 0.111 | 0.79 | 0.57 | 0.79 | 0.58 | 0.75 | 0.66 |
| 8 | 9412153 | rs9329203 | 2.453 | 1.446 | 0.347 | 2.325 | 0.037 | 0.142 | 0.142 | 0.060 | 0.81 | 0.63 | 0.80 | 0.57 | 0.77 | 0.67 |
| 8 | 9540693 | rs13273033 | 2.063 | 1.162 | 0.253 | 2.506 | 0.018 | 0.055 | 0.055 | 0.062 | 0.83 | 0.72 | 0.82 | 0.67 | 0.78 | 0.70 |
| 8 | 9555092 | rs4841196 | 2.858 | 1.204 | 0.344 | 2.541 | 0.006 | 0.046 | 0.046 | 1.579 | 0.74 | 0.61 | 0.57 | 0.38 | 0.72 | 0.57 |
| 8 | 9578982 | rs13276086 | 3.030 | 1.239 | 0.388 | 2.146 | 0.020 | 0.076 | 0.076 | 2.172 | 0.72 | 0.55 | 0.54 | 0.46 | 0.67 | 0.57 |
| 8 | 9601699 | rs12545912 | 2.337 | 0.896 | 0.475 | 2.225 | 0.017 | 0.043 | 0.043 | 1.002 | 0.80 | 0.65 | 0.84 | 0.72 | 0.77 | 0.69 |
| 8 | 17630019 | rs6998733 | 2.191 | 0.073 | 1.727 | 1.459 | 0.018 | 0.080 | 0.080 | 0.137 | 0.50 | 0.33 | 0.47 | 0.16 | 0.51 | 0.18 |
| 8 | 18665597 | rs11778064 | 2.347 | 1.025 | 0.394 | 1.944 | 0.002 | 0.030 | 0.030 | 0.136 | 0.57 | 0.22 | 0.42 | 0.36 | 0.60 | 0.35 |
| 8 | 24144598 | rs436760 | 2.026 | 1.255 | 2.085 | 3.247 | 0.021 | 0.077 | 0.077 | 1.680 | 0.76 | 0.44 | 0.76 | 0.45 | 0.70 | 0.62 |
| 8 | 29957816 | rs12386822 | 2.480 | 0.192 | 4.100 | 3.719 | 0.011 | 0.016 | 0.016 | 0.147 | 0.64 | 0.26 | 0.70 | 0.41 | 0.68 | 0.56 |
| 8 | 29996824 | rs7838088 | 2.706 | 0.284 | 3.204 | 3.442 | 0.006 | 0.241 | 0.241 | 0.380 | 0.88 | 0.56 | 0.84 | 0.71 | 0.91 | 0.93 |
| 8 | 30000346 | rs12334730 | 2.948 | 0.263 | 3.204 | 3.411 | 0.028 | 0.305 | 0.305 | 1.792 | 0.92 | 0.46 | 0.86 | 0.72 | 0.94 | 0.93 |
| 8 | 30004493 | rs2341177 | 2.517 | 0.264 | 3.245 | 3.427 | 0.007 | 0.120 | 0.120 | 0.175 | 0.89 | 0.64 | 0.89 | 0.77 | 0.92 | 0.94 |
| 8 | 33853868 | rs7465297 | 2.173 | 0.865 | 1.521 | 0.466 | 0.007 | 0.017 | 0.017 | 0.723 | 0.78 | 0.31 | 0.76 | 0.71 | 0.79 | 0.80 |
| 8 | 108522424 | rs7003565 | 2.443 | 1.019 | 0.247 | 0.070 | 0.015 | 0.136 | 0.136 | 0.018 | 0.49 | 0.18 | 0.32 | 0.30 | 0.49 | 0.43 |
| 8 | 116701130 | rs12682033 | 2.987 | 1.262 | 3.240 | 1.250 | 0.099 | 0.202 | 0.202 | 0.122 | 0.85 | 0.49 | 0.67 | 0.64 | 0.79 | 0.78 |
| 8 | 139628889 | rs4076991 | 3.181 | 1.091 | 4.328 | 1.093 | 0.030 | 0.128 | 0.128 | 0.285 | 0.87 | 0.51 | 0.76 | 0.79 | 0.89 | 0.86 |
| 8 | 139669119 | rs6989017 | 2.454 | 0.910 | 4.634 | 1.065 | 0.004 | 0.103 | 0.103 | 0.645 | 0.84 | 0.48 | 0.69 | 0.76 | 0.90 | 0.77 |
| 8 | 139751900 | rs10109866 | 2.167 | 3.109 | 3.397 | 1.282 | 0.016 | 0.005 | 0.005 | 0.823 | 0.81 | 0.27 | 0.80 | 0.76 | 0.79 | 0.85 |
| 8 | 139944993 | rs11166859 | 2.814 | 1.316 | 2.820 | 0.246 | 0.089 | 0.030 | 0.030 | 0.306 | 0.91 | 0.61 | 0.86 | 0.95 | 0.90 | 0.81 |
| 8 | 140006609 | rs2318832 | 2.019 | 0.575 | 1.431 | 1.345 | 0.029 | 0.036 | 0.036 | 0.655 | 0.74 | 0.20 | 0.69 | 0.81 | 0.87 | 0.76 |
| 8 | 140031004 | rs10095842 | 2.314 | 1.051 | 3.303 | 2.403 | 0.001 | 0.010 | 0.010 | 0.363 | 0.80 | 0.81 | 0.71 | 0.55 | 0.83 | 0.71 |
| 9 | 907561 | rs4742545 | 2.144 | 1.255 | 2.052 | 1.162 | 0.175 | 0.109 | 0.109 | 0.696 | 0.93 | 0.68 | 0.88 | 0.90 | 0.87 | 0.91 |
| 9 | 14535957 | rs10810175 | 2.025 | 2.896 | 0.805 | 0.988 | 0.080 | 0.129 | 0.129 | 0.001 | 0.45 | 0.25 | 0.30 | 0.17 | 0.29 | 0.28 |
| 9 | 27869510 | rs2383733 | 2.096 | 1.062 | 1.665 | 1.762 | 0.021 | 0.364 | 0.364 | 0.730 | 0.90 | 0.58 | 0.74 | 0.65 | 0.93 | 0.88 |
| 9 | 32698601 | rs1329005 | 2.639 | 0.738 | 3.266 | 0.719 | 0.034 | 0.396 | 0.396 | 4.659 | 0.94 | 0.62 | 0.90 | 0.76 | 1.00 | 0.96 |
| 9 | 32994500 | rs2183870 | 2.029 | 0.764 | 0.636 | 0.790 | 0.005 | 0.006 | 0.006 | 0.472 | 0.47 | 0.31 | 0.53 | 0.44 | 0.49 | 0.57 |
| 9 | 79404749 | rs7854027 | 2.308 | 0.923 | 0.199 | 0.445 | 0.017 | 0.022 | 0.022 | 1.409 | 0.35 | 0.27 | 0.28 | 0.21 | 0.25 | 0.23 |
| 9 | 87995971 | rs2814737 | 2.159 | 0.009 | 1.296 | 0.542 | 0.006 | 0.049 | 0.049 | 0.288 | 0.58 | 0.35 | 0.44 | 0.22 | 0.62 | 0.62 |
| 9 | 90821421 | rs2482488 | 3.205 | 0.973 | 3.271 | 3.037 | 0.058 | 0.209 | 0.209 | 0.013 | 0.81 | 0.57 | 0.63 | 0.48 | 0.80 | 0.69 |
| 9 | 90835589 | rs2578246 | 2.493 | 1.052 | 2.720 | 2.953 | 0.057 | 0.153 | 0.153 | 1.990 | 0.84 | 0.41 | 0.72 | 0.63 | 0.82 | 0.83 |
| 9 | 114731867 | rs10733581 | 2.485 | 0.085 | 1.809 | 2.540 | 0.029 | 0.064 | 0.064 | 0.061 | 0.73 | 0.53 | 0.52 | 0.65 | 0.67 | 0.71 |
| 9 | 123480866 | rs2416799 | 2.329 | 1.074 | 2.000 | 0.735 | 0.003 | 0.024 | 0.024 | 0.926 | 0.68 | 0.38 | 0.75 | 0.58 | 0.70 | 0.74 |
| 10 | 30937956 | rs2015920 | 2.059 | 1.282 | 1.801 | 0.949 | 0.019 | 0.021 | 0.021 | 1.647 | 0.75 | 0.42 | 0.61 | 0.67 | 0.73 | 0.62 |
| 10 | 30985081 | rs10826846 | 2.168 | 0.774 | 1.915 | 2.007 | 0.005 | 0.251 | 0.251 | 4.651 | 0.89 | 0.38 | 0.85 | 0.66 | 0.91 | 0.97 |
| 10 | 44438863 | rs11238782 | 2.314 | 0.069 | 0.593 | 1.416 | 0.001 | 0.047 | 0.047 | 2.957 | 0.53 | 0.15 | 0.33 | 0.40 | 0.59 | 0.47 |
| 10 | 78314705 | rs2637256 | 2.245 | 0.626 | 2.643 | 1.783 | 0.002 | 0.496 | 0.496 | 0.493 | 0.90 | 0.52 | 0.85 | 0.52 | 0.96 | 0.85 |
| 10 | 78316518 | rs2579766 | 2.286 | 0.653 | 2.688 | 1.791 | 0.006 | 0.504 | 0.504 | 0.418 | 0.90 | 0.52 | 0.85 | 0.52 | 0.96 | 0.85 |
| 10 | 80569876 | rs7477887 | 2.207 | 1.669 | 0.814 | 0.676 | 0.011 | 0.062 | 0.062 | 0.104 | 0.46 | 0.22 | 0.44 | 0.36 | 0.41 | 0.43 |
| 10 | 90033593 | rs2114406 | 2.121 | 0.546 | 0.660 | 1.029 | 0.062 | 0.099 | 0.099 | 1.952 | 0.42 | 0.14 | 0.30 | 0.23 | 0.36 | 0.39 |
| 10 | 108260332 | rs2900712 | 2.686 | 2.217 | 0.313 | 1.647 | 0.031 | 0.016 | 0.016 | 0.784 | 0.51 | 0.21 | 0.30 | 0.44 | 0.42 | 0.38 |
| 10 | 108495751 | rs2756241 | 2.600 | 0.645 | 1.545 | 2.621 | 0.008 | 0.282 | 0.282 | 1.565 | 0.85 | 0.58 | 0.80 | 0.52 | 0.88 | 0.72 |
| 10 | 108495847 | rs2756242 | 2.600 | 0.645 | 1.547 | 2.621 | 0.008 | 0.286 | 0.286 | 1.565 | 0.85 | 0.56 | 0.80 | 0.52 | 0.88 | 0.72 |
| 10 | 119994707 | rs10787813 | 2.099 | 0.188 | 1.038 | 0.404 | 0.017 | 0.012 | 0.012 | 0.644 | 0.89 | 0.44 | 0.90 | 0.93 | 0.91 | 0.91 |
| 10 | 120828969 | rs7908387 | 2.071 | 0.602 | 1.377 | 0.860 | 0.010 | 0.141 | 0.141 | 4.168 | 0.92 | 0.39 | 0.92 | 0.88 | 0.97 | 0.96 |
| 10 | 121932833 | rs4427485 | 2.187 | 0.463 | 2.207 | 0.037 | 0.006 | 0.043 | 0.043 | 0.185 | 0.49 | 0.24 | 0.60 | 0.60 | 0.52 | 0.51 |
| 10 | 128837927 | rs9418809 | 2.168 | 0.468 | 0.243 | 0.351 | 0.013 | 0.020 | 0.020 | 0.014 | 0.27 | 0.22 | 0.23 | 0.21 | 0.22 | 0.22 |
| 10 | 131236879 | rs10734086 | 2.671 | 1.067 | 2.721 | 0.213 | 0.010 | 0.107 | 0.107 | 0.664 | 0.86 | 0.48 | 0.79 | 0.77 | 0.88 | 0.80 |
| 10 | 133533485 | rs4367873 | 2.446 | 0.393 | 1.777 | 0.810 | 0.004 | 0.007 | 0.007 | 0.130 | 0.70 | 0.23 | 0.76 | 0.78 | 0.66 | 0.79 |
| 10 | 133534822 | rs9419624 | 2.268 | 0.374 | 1.750 | 0.784 | 0.005 | 0.011 | 0.011 | 0.089 | 0.69 | 0.22 | 0.75 | 0.78 | 0.65 | 0.77 |
| 11 | 15580347 | rs2351046 | 2.350 | 3.135 | 1.609 | 0.648 | 0.014 | 0.009 | 0.009 | 0.028 | 0.67 | 0.38 | 0.54 | 0.55 | 0.56 | 0.60 |
| 11 | 15610823 | rs4756814 | 2.627 | 2.896 | 1.707 | 0.669 | 0.023 | 0.015 | 0.015 | 0.160 | 0.72 | 0.36 | 0.65 | 0.58 | 0.62 | 0.66 |
| 11 | 15616694 | rs11023650 | 2.130 | 2.690 | 1.605 | 0.670 | 0.029 | 0.016 | 0.016 | 0.139 | 0.70 | 0.39 | 0.63 | 0.58 | 0.59 | 0.64 |
| 11 | 20483748 | rs3758803 | 2.053 | 0.261 | 1.161 | 1.464 | 0.041 | 0.129 | 0.129 | 0.301 | 0.50 | 0.19 | 0.39 | 0.22 | 0.57 | 0.42 |
| 11 | 20776391 | rs1554367 | 2.409 | 0.827 | 2.432 | 1.091 | 0.093 | 0.014 | 0.014 | 1.480 | 0.87 | 0.44 | 0.82 | 0.88 | 0.82 | 0.85 |

| CHR | POS | SNP | iHS | XP-EHH | | | PBS | | | LLRS | Alternate Allele Frequency | | | | | |
|---|---|---|---|---|---|---|---|---|---|---|---|---|---|---|---|---|
| | | | | KWT vs CEU | KWT vs YRI | KWT vs CHB | KWT vs CEU | KWT vs YRI | KWT vs CHB | | KWT | AFR | AMR | EAS | EUR | SAS |
| 11 | 22696451 | rs922571 | 2.331 | 0.234 | 0.531 | 0.428 | 0.012 | 0.055 | 0.055 | 0.471 | 0.64 | 0.17 | 0.61 | 0.75 | 0.67 | 0.55 |
| 11 | 34283861 | rs2611145 | 2.022 | 0.005 | 0.414 | 1.359 | 0.014 | 0.034 | 0.034 | 0.301 | 0.53 | 0.33 | 0.46 | 0.39 | 0.53 | 0.62 |
| 11 | 45631953 | rs4755330 | 2.119 | 1.582 | 1.613 | 0.180 | 0.120 | 0.017 | 0.017 | 0.311 | 0.85 | 0.47 | 0.83 | 0.87 | 0.78 | 0.85 |
| 11 | 45672261 | rs3802762 | 2.754 | 0.977 | 1.445 | 0.927 | 0.076 | 0.194 | 0.194 | 0.579 | 0.88 | 0.40 | 0.78 | 0.67 | 0.83 | 0.88 |
| 11 | 45672725 | rs4148903 | 2.516 | 0.976 | 1.577 | 0.930 | 0.134 | 0.288 | 0.288 | 0.795 | 0.89 | 0.47 | 0.78 | 0.68 | 0.83 | 0.89 |
| 11 | 45672747 | rs4148902 | 3.144 | 0.975 | 1.528 | 0.915 | 0.067 | 0.180 | 0.180 | 1.470 | 0.87 | 0.39 | 0.78 | 0.67 | 0.83 | 0.88 |
| 11 | 45674170 | rs2277272 | 3.369 | 0.957 | 1.489 | 0.879 | 0.052 | 0.149 | 0.149 | 1.155 | 0.86 | 0.39 | 0.77 | 0.67 | 0.82 | 0.87 |
| 11 | 45674599 | rs4148898 | 3.090 | 0.958 | 1.490 | 0.880 | 0.064 | 0.173 | 0.173 | 0.561 | 0.87 | 0.39 | 0.78 | 0.67 | 0.83 | 0.88 |
| 11 | 56832642 | rs550124 | 2.052 | 2.534 | 1.156 | 1.412 | 0.039 | 0.098 | 0.098 | 8.279 | 0.42 | 0.13 | 0.45 | 0.32 | 0.39 | 0.42 |
| 11 | 56952427 | rs10896578 | 2.555 | 1.997 | 0.974 | 0.674 | 0.010 | 0.003 | 0.003 | 1.568 | 0.43 | 0.12 | 0.43 | 0.36 | 0.40 | 0.52 |
| 11 | 59210550 | rs7947825 | 2.161 | 1.925 | 0.802 | 0.875 | 0.061 | 0.160 | 0.160 | 1.421 | 0.91 | 0.48 | 0.72 | 0.87 | 0.95 | 0.89 |
| 11 | 67068859 | rs2298815 | 2.194 | 0.727 | 3.371 | 0.717 | 0.002 | 0.063 | 0.063 | 0.015 | 0.94 | 0.76 | 0.95 | 0.89 | 0.96 | 0.98 |
| 11 | 67070017 | rs4930192 | 3.309 | 0.727 | 3.551 | 0.718 | 0.040 | 0.173 | 0.173 | 0.316 | 0.90 | 0.54 | 0.88 | 0.78 | 0.93 | 0.95 |
| 11 | 67419855 | rs10736660 | 2.330 | 0.862 | 2.182 | 0.795 | 0.014 | 0.040 | 0.040 | 0.755 | 0.82 | 0.43 | 0.85 | 0.73 | 0.81 | 0.92 |
| 11 | 76814110 | rs2233549 | 2.020 | 0.237 | 0.135 | 0.375 | 0.005 | 0.012 | 0.012 | 0.192 | 0.18 | 0.20 | 0.21 | 0.23 | 0.21 | 0.36 |
| 11 | 83744306 | rs10792710 | 3.236 | 0.496 | 3.111 | 0.820 | 0.009 | 0.002 | 0.002 | 0.242 | 0.72 | 0.24 | 0.68 | 0.77 | 0.75 | 0.67 |
| 11 | 83766621 | rs4943885 | 2.290 | 0.648 | 2.467 | 0.924 | 0.025 | 0.107 | 0.107 | 0.013 | 0.84 | 0.50 | 0.74 | 0.77 | 0.83 | 0.81 |
| 11 | 83839293 | rs2105806 | 3.207 | 1.175 | 3.470 | 1.371 | 0.001 | 0.011 | 0.011 | 0.265 | 0.80 | 0.30 | 0.69 | 0.74 | 0.78 | 0.77 |
| 11 | 83844790 | rs1945825 | 3.338 | 1.167 | 3.403 | 1.388 | 0.014 | 0.051 | 0.051 | 0.078 | 0.80 | 0.43 | 0.70 | 0.74 | 0.78 | 0.77 |
| 11 | 83904512 | rs1349820 | 3.075 | 1.032 | 2.708 | 1.361 | 0.000 | 0.018 | 0.018 | 0.351 | 0.78 | 0.35 | 0.69 | 0.75 | 0.78 | 0.78 |
| 11 | 83918339 | rs10792731 | 2.941 | 1.002 | 2.699 | 1.391 | 0.016 | 0.115 | 0.115 | 0.063 | 0.85 | 0.40 | 0.73 | 0.75 | 0.85 | 0.79 |
| 11 | 83933781 | rs7925243 | 3.039 | 1.139 | 2.766 | 1.332 | 0.018 | 0.053 | 0.053 | 0.232 | 0.79 | 0.40 | 0.70 | 0.74 | 0.77 | 0.78 |
| 11 | 83939591 | rs10792740 | 3.169 | 1.127 | 2.898 | 1.338 | 0.047 | 0.197 | 0.197 | 0.199 | 0.86 | 0.50 | 0.73 | 0.74 | 0.84 | 0.83 |
| 11 | 83945482 | rs896996 | 3.208 | 1.133 | 2.941 | 1.331 | 0.028 | 0.107 | 0.107 | 0.134 | 0.81 | 0.46 | 0.71 | 0.74 | 0.77 | 0.80 |
| 11 | 83954724 | rs1399617 | 3.456 | 1.077 | 2.817 | 1.352 | 0.020 | 0.087 | 0.087 | 0.067 | 0.79 | 0.49 | 0.70 | 0.74 | 0.75 | 0.76 |
| 11 | 83967019 | rs4943889 | 3.318 | 0.975 | 2.625 | 1.469 | 0.016 | 0.116 | 0.116 | 0.464 | 0.80 | 0.57 | 0.70 | 0.73 | 0.78 | 0.84 |
| 11 | 83976634 | rs10792746 | 2.199 | 1.059 | 2.111 | 1.429 | 0.043 | 0.103 | 0.103 | 0.140 | 0.80 | 0.55 | 0.71 | 0.75 | 0.78 | 0.86 |
| 11 | 84993643 | rs552017 | 2.070 | 0.823 | 0.403 | 0.020 | 0.010 | 0.164 | 0.164 | 0.091 | 0.85 | 0.62 | 0.62 | 0.60 | 0.85 | 0.63 |
| 11 | 88452034 | rs1993842 | 2.190 | 0.690 | 1.765 | 1.153 | 0.100 | 0.274 | 0.274 | 1.141 | 0.93 | 0.48 | 0.76 | 0.83 | 0.95 | 0.94 |
| 11 | 102495066 | rs1962082 | 2.121 | 3.033 | 0.117 | 0.459 | 0.045 | 0.073 | 0.073 | 0.134 | 0.42 | 0.27 | 0.26 | 0.05 | 0.28 | 0.33 |
| 11 | 104758035 | rs484438 | 2.490 | 1.116 | 2.959 | 1.113 | 0.052 | 0.054 | 0.054 | 2.077 | 0.95 | 0.62 | 0.96 | 1.00 | 0.99 | 0.98 |
| 11 | 107987784 | rs10890813 | 2.195 | 2.648 | 0.411 | 0.732 | 0.138 | 0.059 | 0.059 | 1.394 | 0.50 | 0.06 | 0.42 | 0.34 | 0.34 | 0.51 |
| 11 | 108038477 | rs920679 | 2.095 | 2.773 | 1.121 | 0.976 | 0.125 | 0.033 | 0.033 | 0.635 | 0.49 | 0.06 | 0.45 | 0.36 | 0.36 | 0.52 |
| 11 | 128818599 | rs4121427 | 2.185 | 2.498 | 0.874 | 1.579 | 0.030 | 0.003 | 0.003 | 0.000 | 0.42 | 0.26 | 0.46 | 0.41 | 0.44 | 0.25 |
| 11 | 128823766 | rs4121417 | 2.830 | 2.357 | 0.831 | 1.575 | 0.054 | 0.019 | 0.019 | 0.220 | 0.36 | 0.20 | 0.39 | 0.32 | 0.32 | 0.14 |
| 11 | 128843573 | rs10893942 | 3.251 | 2.379 | 1.146 | 1.724 | 0.041 | 0.031 | 0.031 | 0.058 | 0.35 | 0.17 | 0.37 | 0.27 | 0.32 | 0.13 |
| 11 | 128857025 | rs11221527 | 3.253 | 2.366 | 1.148 | 1.691 | 0.041 | 0.021 | 0.021 | 0.288 | 0.35 | 0.17 | 0.37 | 0.30 | 0.32 | 0.13 |
| 11 | 128927178 | rs10893962 | 2.866 | 2.284 | 1.034 | 1.685 | 0.044 | 0.020 | 0.020 | 0.272 | 0.35 | 0.17 | 0.37 | 0.30 | 0.31 | 0.14 |
| 11 | 128937691 | rs2840345 | 2.827 | 2.228 | 1.035 | 1.679 | 0.044 | 0.020 | 0.020 | 0.272 | 0.35 | 0.17 | 0.37 | 0.30 | 0.31 | 0.14 |
| 11 | 129103092 | rs10219187 | 2.598 | 2.024 | 0.999 | 1.621 | 0.031 | 0.029 | 0.029 | 0.025 | 0.36 | 0.26 | 0.37 | 0.27 | 0.32 | 0.18 |
| 11 | 129126706 | rs4372462 | 2.664 | 2.079 | 1.068 | 1.146 | 0.010 | 0.011 | 0.011 | 0.076 | 0.36 | 0.31 | 0.38 | 0.19 | 0.32 | 0.15 |
| 11 | 129150067 | rs4077032 | 2.518 | 1.949 | 0.866 | 1.009 | 0.031 | 0.057 | 0.057 | 0.169 | 0.35 | 0.23 | 0.38 | 0.22 | 0.32 | 0.15 |
| 11 | 130060217 | rs478745 | 2.380 | 0.515 | 0.541 | 1.675 | 0.006 | 0.002 | 0.002 | 0.838 | 0.67 | 0.12 | 0.75 | 0.71 | 0.73 | 0.83 |
| 12 | 4915652 | rs12821926 | 2.656 | 1.842 | 0.652 | 0.836 | 0.015 | 0.041 | 0.041 | 3.890 | 0.35 | 0.22 | 0.18 | 0.03 | 0.31 | 0.06 |
| 12 | 11156123 | rs10772413 | 2.006 | 1.505 | 1.485 | 0.031 | 0.037 | 0.134 | 0.134 | 0.341 | 0.55 | 0.06 | 0.38 | 0.76 | 0.38 | 0.63 |
| 12 | 11164102 | rs7304447 | 2.004 | 1.518 | 1.491 | 0.037 | 0.037 | 0.134 | 0.134 | 0.349 | 0.55 | 0.07 | 0.38 | 0.76 | 0.38 | 0.63 |
| 12 | 11827629 | rs2111049 | 2.511 | 0.230 | 0.679 | 1.300 | 0.036 | 0.003 | 0.003 | 0.779 | 0.75 | 0.11 | 0.80 | 0.82 | 0.87 | 0.84 |
| 12 | 11830715 | rs2724626 | 2.512 | 0.218 | 0.597 | 1.291 | 0.037 | 0.006 | 0.006 | 0.850 | 0.75 | 0.11 | 0.80 | 0.82 | 0.86 | 0.84 |
| 12 | 76327701 | rs6582316 | 2.130 | 0.579 | 0.758 | 2.664 | 0.077 | 0.217 | 0.217 | 0.667 | 0.91 | 0.75 | 0.89 | 0.65 | 0.89 | 0.65 |
| 12 | 81522025 | rs2574740 | 2.186 | 0.494 | 1.690 | 2.176 | 0.096 | 0.463 | 0.463 | 3.345 | 0.93 | 0.53 | 0.93 | 0.69 | 0.96 | 0.81 |
| 12 | 83361414 | rs10862531 | 2.127 | 0.769 | 1.788 | 0.489 | 0.016 | 0.053 | 0.053 | 3.191 | 0.81 | 0.38 | 0.77 | 0.71 | 0.79 | 0.88 |
| 12 | 83419504 | rs11115544 | 2.547 | 0.713 | 2.214 | 0.492 | 0.003 | 0.014 | 0.014 | 1.832 | 0.67 | 0.39 | 0.63 | 0.57 | 0.65 | 0.75 |
| 12 | 104943735 | rs17207200 | 2.182 | 1.863 | 0.413 | 0.335 | 0.094 | 0.202 | 0.202 | 1.473 | 0.44 | 0.14 | 0.31 | 0.19 | 0.28 | 0.40 |
| 12 | 104962244 | rs313318 | 2.110 | 1.885 | 0.002 | 0.381 | 0.065 | 0.166 | 0.166 | 1.312 | 0.42 | 0.12 | 0.32 | 0.19 | 0.31 | 0.37 |
| 12 | 109633316 | rs4300440 | 2.089 | 1.792 | 1.487 | 1.964 | 0.020 | 0.002 | 0.002 | 0.585 | 0.67 | 0.63 | 0.64 | 0.32 | 0.60 | 0.49 |
| 12 | 121359586 | rs4767941 | 2.107 | 2.566 | 0.071 | 0.374 | 0.086 | 0.011 | 0.011 | 1.084 | 0.51 | 0.07 | 0.54 | 0.43 | 0.38 | 0.41 |

Table S2. The selection tests scores and the allele frequency of 385 variants - Continued

| CHR | POS | SNP | iHS | XP-EHH | | | PBS | | | LLRS | Alternate Allele Frequency | | | | | |
|---|---|---|---|---|---|---|---|---|---|---|---|---|---|---|---|---|
| | | | | KWT vs CEU | KWT vs YRI | KWT vs CHB | KWT vs CEU | KWT vs YRI | KWT vs CHB | | KWT | AFR | AMR | EAS | EUR | SAS |
| 12 | 121476423 | rs10774580 | 2.099 | 2.551 | 1.143 | 0.388 | 0.000 | 0.000 | 0.000 | 0.333 | 0.65 | 0.34 | 0.70 | 0.58 | 0.62 | 0.76 |
| 12 | 125125778 | rs2088229 | 2.143 | 2.967 | 1.829 | 1.227 | 0.089 | 0.015 | 0.015 | 0.097 | 0.81 | 0.43 | 0.71 | 0.78 | 0.69 | 0.78 |
| 12 | 125125932 | rs188025 | 2.121 | 2.964 | 1.816 | 1.219 | 0.071 | 0.015 | 0.015 | 0.078 | 0.81 | 0.41 | 0.75 | 0.78 | 0.73 | 0.79 |
| 12 | 125134746 | rs4542507 | 2.604 | 1.814 | 0.873 | 0.963 | 0.032 | 0.129 | 0.129 | 0.018 | 0.45 | 0.16 | 0.35 | 0.22 | 0.39 | 0.48 |
| 13 | 21510900 | rs4770084 | 2.087 | 0.612 | 1.694 | 1.067 | 0.025 | 0.287 | 0.287 | 2.723 | 0.88 | 0.45 | 0.86 | 0.63 | 0.95 | 0.79 |
| 13 | 22652247 | rs1323172 | 2.291 | 0.546 | 1.593 | 0.716 | 0.042 | 0.033 | 0.033 | 4.758 | 0.94 | 0.38 | 0.95 | 0.99 | 1.00 | 0.91 |
| 13 | 46093829 | rs2985980 | 2.216 | 2.468 | 1.643 | 0.982 | 0.005 | 0.008 | 0.008 | 2.160 | 0.82 | 0.31 | 0.83 | 0.78 | 0.79 | 0.77 |
| 13 | 46098994 | rs3014966 | 2.087 | 2.646 | 1.580 | 0.974 | 0.048 | 0.031 | 0.031 | 1.245 | 0.82 | 0.44 | 0.81 | 0.78 | 0.74 | 0.76 |
| 13 | 46103935 | rs2274285 | 2.087 | 2.643 | 1.302 | 0.975 | 0.048 | 0.031 | 0.031 | 1.245 | 0.82 | 0.44 | 0.82 | 0.78 | 0.74 | 0.76 |
| 13 | 67967187 | rs9541094 | 2.583 | 1.641 | 2.134 | 0.880 | 0.095 | 0.023 | 0.023 | 0.172 | 0.91 | 0.58 | 0.88 | 0.95 | 0.93 | 0.96 |
| 13 | 72026200 | rs2325392 | 2.131 | 0.436 | 1.434 | 1.055 | 0.044 | 0.075 | 0.075 | 3.776 | 0.94 | 0.49 | 0.93 | 0.97 | 0.99 | 0.99 |
| 13 | 72510750 | rs9542784 | 2.424 | 0.850 | 3.062 | 1.526 | 0.075 | 0.085 | 0.085 | 0.751 | 0.90 | 0.56 | 0.90 | 0.88 | 0.92 | 0.94 |
| 13 | 72598561 | rs2810131 | 2.071 | 1.167 | 3.776 | 0.876 | 0.017 | 0.011 | 0.011 | 0.024 | 0.85 | 0.68 | 0.90 | 0.82 | 0.87 | 0.81 |
| 13 | 72599382 | rs2706420 | 3.000 | 1.192 | 3.850 | 0.907 | 0.100 | 0.042 | 0.042 | 0.105 | 0.86 | 0.52 | 0.77 | 0.81 | 0.80 | 0.72 |
| 13 | 83226757 | rs9318893 | 2.527 | 2.220 | 0.929 | 1.282 | 0.039 | 0.088 | 0.088 | 0.049 | 0.51 | 0.23 | 0.47 | 0.31 | 0.39 | 0.45 |
| 13 | 93831943 | rs319561 | 2.716 | 1.271 | 1.992 | 1.570 | 0.014 | 0.010 | 0.010 | 0.067 | 0.77 | 0.35 | 0.62 | 0.65 | 0.75 | 0.71 |
| 13 | 108360217 | rs4113422 | 2.857 | 0.972 | 2.582 | 2.578 | 0.045 | 0.358 | 0.358 | 0.105 | 0.92 | 0.71 | 0.88 | 0.58 | 0.92 | 0.87 |
| 13 | 108383142 | rs3905064 | 2.549 | 1.167 | 1.738 | 2.347 | 0.032 | 0.276 | 0.276 | 2.060 | 0.84 | 0.50 | 0.80 | 0.54 | 0.84 | 0.73 |
| 13 | 108385281 | rs7330529 | 2.357 | 1.132 | 1.687 | 2.351 | 0.013 | 0.303 | 0.303 | 1.288 | 0.86 | 0.57 | 0.82 | 0.54 | 0.88 | 0.74 |
| 14 | 63555491 | rs2146940 | 3.210 | 0.191 | 0.854 | 2.350 | 0.010 | 0.103 | 0.103 | 1.501 | 0.57 | 0.07 | 0.57 | 0.29 | 0.65 | 0.68 |
| 14 | 63568154 | rs1951805 | 2.856 | 0.545 | 1.070 | 2.335 | 0.009 | 0.012 | 0.012 | 3.795 | 0.61 | 0.37 | 0.66 | 0.33 | 0.60 | 0.63 |
| 14 | 63572267 | rs10129357 | 2.275 | 0.424 | 1.466 | 2.347 | 0.029 | 0.109 | 0.109 | 0.811 | 0.89 | 0.63 | 0.89 | 0.80 | 0.87 | 0.94 |
| 14 | 63691272 | rs10150375 | 3.126 | 1.331 | 3.908 | 3.914 | 0.048 | 0.289 | 0.289 | 0.764 | 0.92 | 0.61 | 0.90 | 0.70 | 0.91 | 0.94 |
| 14 | 85548835 | rs899760 | 2.154 | 0.780 | 1.212 | 0.011 | 0.014 | 0.011 | 0.011 | 1.790 | 0.67 | 0.29 | 0.80 | 0.69 | 0.74 | 0.65 |
| 14 | 89430335 | rs1885185 | 2.282 | 1.544 | 0.278 | 0.409 | 0.030 | 0.059 | 0.059 | 1.445 | 0.35 | 0.19 | 0.20 | 0.19 | 0.26 | 0.18 |
| 14 | 89433411 | rs17702956 | 2.236 | 1.370 | 0.309 | 0.434 | 0.038 | 0.100 | 0.100 | 1.552 | 0.35 | 0.09 | 0.19 | 0.19 | 0.26 | 0.18 |
| 14 | 100993962 | rs4905969 | 2.454 | 1.318 | 2.024 | 0.471 | 0.027 | 0.133 | 0.133 | 4.269 | 0.93 | 0.46 | 0.96 | 0.89 | 1.00 | 0.96 |
| 15 | 61892544 | rs12899273 | 2.614 | 0.294 | 1.121 | 0.368 | 0.025 | 0.050 | 0.050 | 2.565 | 0.67 | 0.22 | 0.78 | 0.84 | 0.75 | 0.63 |
| 15 | 61901612 | rs11634723 | 2.438 | 0.416 | 0.863 | 0.385 | 0.021 | 0.053 | 0.053 | 3.012 | 0.66 | 0.21 | 0.77 | 0.81 | 0.71 | 0.60 |
| 15 | 68615873 | rs1516869 | 2.720 | 1.380 | 1.401 | 1.025 | 0.005 | 0.029 | 0.029 | 0.099 | 0.73 | 0.21 | 0.81 | 0.80 | 0.78 | 0.79 |
| 15 | 68626542 | rs964691 | 2.527 | 2.453 | 1.190 | 0.990 | 0.004 | 0.018 | 0.018 | 0.534 | 0.75 | 0.22 | 0.82 | 0.80 | 0.79 | 0.79 |
| 15 | 68628143 | rs2306023 | 2.325 | 2.464 | 1.664 | 1.003 | 0.003 | 0.014 | 0.014 | 0.368 | 0.75 | 0.23 | 0.82 | 0.80 | 0.79 | 0.78 |
| 15 | 70951141 | rs7174472 | 2.336 | 1.019 | 2.368 | 0.810 | 0.005 | 0.071 | 0.071 | 0.072 | 0.76 | 0.41 | 0.82 | 0.54 | 0.77 | 0.65 |
| 15 | 70960068 | rs934005 | 2.243 | 1.037 | 2.276 | 0.809 | 0.000 | 0.016 | 0.016 | 0.479 | 0.75 | 0.32 | 0.74 | 0.53 | 0.76 | 0.64 |
| 15 | 72613485 | rs11072355 | 2.073 | 0.017 | 2.934 | 0.043 | 0.005 | 0.131 | 0.131 | 3.421 | 0.92 | 0.66 | 0.96 | 0.79 | 0.97 | 0.96 |
| 15 | 73235674 | rs1823338 | 2.089 | 0.192 | 2.034 | 1.028 | 0.002 | 0.023 | 0.023 | 2.444 | 0.65 | 0.42 | 0.59 | 0.55 | 0.65 | 0.57 |
| 15 | 73259994 | rs921261 | 2.132 | 0.188 | 2.241 | 1.004 | 0.009 | 0.044 | 0.044 | 1.764 | 0.66 | 0.51 | 0.60 | 0.54 | 0.65 | 0.59 |
| 15 | 95782720 | rs7166820 | 2.792 | 1.586 | 2.654 | 1.055 | 0.006 | 0.003 | 0.003 | 0.571 | 0.64 | 0.45 | 0.62 | 0.68 | 0.58 | 0.55 |
| 15 | 97251036 | rs4567667 | 2.010 | 1.447 | 0.289 | 0.628 | 0.008 | 0.089 | 0.089 | 0.173 | 0.83 | 0.55 | 0.87 | 0.69 | 0.85 | 0.76 |
| 15 | 99656047 | rs1965866 | 2.391 | 2.095 | 1.098 | 0.806 | 0.042 | 0.037 | 0.037 | 4.381 | 0.51 | 0.23 | 0.40 | 0.35 | 0.40 | 0.36 |
| 15 | 99663230 | rs7162579 | 2.508 | 2.285 | 1.294 | 0.881 | 0.036 | 0.043 | 0.043 | 5.009 | 0.50 | 0.29 | 0.40 | 0.36 | 0.41 | 0.36 |
| 15 | 99688194 | rs12591477 | 2.893 | 2.139 | 2.078 | 1.244 | 0.006 | 0.032 | 0.032 | 5.709 | 0.50 | 0.19 | 0.42 | 0.36 | 0.44 | 0.35 |
| 15 | 99739889 | rs1377267 | 2.381 | 2.045 | 1.692 | 1.305 | 0.009 | 0.044 | 0.044 | 4.946 | 0.51 | 0.26 | 0.36 | 0.34 | 0.44 | 0.34 |
| 16 | 1873087 | rs104843 | 2.637 | 0.172 | 2.359 | 2.351 | 0.001 | 0.002 | 0.002 | 2.582 | 0.60 | 0.56 | 0.64 | 0.52 | 0.61 | 0.45 |
| 16 | 1901211 | rs7200137 | 2.707 | 0.215 | 2.766 | 2.281 | 0.000 | 0.003 | 0.003 | 2.732 | 0.60 | 0.56 | 0.64 | 0.47 | 0.62 | 0.32 |
| 16 | 1908451 | rs2437744 | 2.015 | 0.243 | 2.286 | 2.295 | 0.003 | 0.014 | 0.014 | 0.686 | 0.71 | 0.64 | 0.68 | 0.58 | 0.69 | 0.64 |
| 16 | 7806373 | rs12445359 | 2.953 | 0.107 | 2.254 | 1.073 | 0.033 | 0.031 | 0.031 | 0.650 | 0.55 | 0.12 | 0.58 | 0.36 | 0.60 | 0.58 |
| 16 | 63096818 | rs8063250 | 2.896 | 1.173 | 0.863 | 0.487 | 0.002 | 0.034 | 0.034 | 0.203 | 0.50 | 0.20 | 0.60 | 0.42 | 0.53 | 0.41 |
| 16 | 63119549 | rs1510213 | 2.539 | 0.948 | 0.653 | 0.504 | 0.019 | 0.021 | 0.021 | 0.249 | 0.52 | 0.22 | 0.64 | 0.41 | 0.61 | 0.44 |
| 16 | 80146783 | rs4889102 | 2.565 | 0.936 | 2.439 | 1.390 | 0.014 | 0.105 | 0.105 | 0.018 | 0.80 | 0.45 | 0.82 | 0.59 | 0.78 | 0.63 |
| 16 | 81070697 | rs2602431 | 3.692 | 0.997 | 1.747 | 1.128 | 0.025 | 0.077 | 0.077 | 0.019 | 0.85 | 0.39 | 0.82 | 0.72 | 0.87 | 0.88 |
| 16 | 81074800 | rs2970077 | 2.702 | 0.852 | 1.859 | 1.061 | 0.054 | 0.006 | 0.006 | 0.014 | 0.89 | 0.43 | 0.93 | 0.92 | 0.91 | 0.98 |
| 16 | 81079641 | rs13185 | 3.638 | 0.840 | 1.811 | 1.091 | 0.011 | 0.079 | 0.079 | 0.053 | 0.85 | 0.38 | 0.83 | 0.72 | 0.88 | 0.88 |
| 16 | 81091257 | rs2911158 | 3.342 | 0.866 | 1.958 | 1.097 | 0.028 | 0.131 | 0.131 | 0.023 | 0.88 | 0.38 | 0.85 | 0.72 | 0.90 | 0.91 |
| 16 | 81122861 | rs8043855 | 2.750 | 0.681 | 1.023 | 0.951 | 0.052 | 0.017 | 0.017 | 0.130 | 0.90 | 0.53 | 0.93 | 0.92 | 0.90 | 0.98 |
| 17 | 53830710 | rs2912547 | 2.679 | 0.311 | 1.490 | 0.823 | 0.046 | 0.125 | 0.125 | 1.280 | 0.68 | 0.07 | 0.70 | 0.86 | 0.80 | 0.80 |

Table S2. The selection tests scores and the allele frequency of 385 variants - Continued

| CHR | POS | SNP | iHS | XP-EHH | | | PBS | | | LLRS | Alternate Allele Frequency | | | | | |
|---|---|---|---|---|---|---|---|---|---|---|---|---|---|---|---|---|
| | | | | KWT vs CEU | KWT vs YRI | KWT vs CHB | KWT vs CEU | KWT vs YRI | KWT vs CHB | | KWT | AFR | AMR | EAS | EUR | SAS |
| 17 | 53840694 | rs2332513 | 2.986 | 0.280 | 1.705 | 0.835 | 0.050 | 0.110 | 0.110 | 1.060 | 0.69 | 0.12 | 0.70 | 0.86 | 0.80 | 0.80 |
| 17 | 53842322 | rs2960070 | 3.066 | 0.280 | 1.698 | 0.835 | 0.046 | 0.125 | 0.125 | 1.230 | 0.68 | 0.07 | 0.70 | 0.86 | 0.80 | 0.80 |
| 17 | 53846348 | rs2960067 | 3.066 | 0.280 | 1.698 | 0.835 | 0.046 | 0.125 | 0.125 | 1.230 | 0.68 | 0.07 | 0.70 | 0.86 | 0.80 | 0.80 |
| 17 | 53852111 | rs2960061 | 2.581 | 0.337 | 1.828 | 0.841 | 0.036 | 0.052 | 0.052 | 0.689 | 0.75 | 0.16 | 0.79 | 0.93 | 0.86 | 0.82 |
| 17 | 53854141 | rs2303506 | 3.031 | 0.332 | 1.771 | 0.834 | 0.039 | 0.128 | 0.128 | 1.401 | 0.68 | 0.07 | 0.69 | 0.86 | 0.80 | 0.80 |
| 17 | 62400000 | rs1050382 | 2.418 | 1.934 | 1.748 | 0.214 | 0.019 | 0.010 | 0.010 | 2.338 | 0.69 | 0.35 | 0.66 | 0.68 | 0.58 | 0.61 |
| 17 | 62401118 | rs2812 | 2.434 | 1.979 | 1.507 | 0.194 | 0.015 | 0.005 | 0.005 | 2.739 | 0.68 | 0.34 | 0.64 | 0.69 | 0.56 | 0.60 |
| 17 | 70968440 | rs4968999 | 2.193 | 0.890 | 1.483 | 1.836 | 0.047 | 0.087 | 0.087 | 0.933 | 0.91 | 0.59 | 0.93 | 0.91 | 0.91 | 0.94 |
| 17 | 71010890 | rs9911924 | 2.336 | 0.693 | 2.082 | 1.108 | 0.031 | 0.045 | 0.045 | 0.912 | 0.90 | 0.41 | 0.83 | 0.86 | 0.92 | 0.90 |
| 17 | 71018801 | rs8069244 | 2.191 | 0.701 | 2.208 | 1.121 | 0.063 | 0.089 | 0.089 | 0.140 | 0.92 | 0.60 | 0.86 | 0.84 | 0.92 | 0.86 |
| 17 | 71019305 | rs4290532 | 2.310 | 0.666 | 2.241 | 1.103 | 0.051 | 0.076 | 0.076 | 0.940 | 0.92 | 0.62 | 0.87 | 0.84 | 0.92 | 0.86 |
| 17 | 77438911 | rs8076334 | 2.137 | 0.765 | 0.297 | 0.266 | 0.014 | 0.036 | 0.036 | 2.154 | 0.54 | 0.34 | 0.28 | 0.28 | 0.42 | 0.40 |
| 18 | 10302548 | rs206457 | 2.511 | 0.009 | 1.240 | 1.378 | 0.013 | 0.020 | 0.020 | 1.303 | 0.49 | 0.21 | 0.42 | 0.43 | 0.52 | 0.38 |
| 18 | 20461282 | rs3863524 | 2.082 | 1.147 | 2.622 | 1.672 | 0.101 | 0.224 | 0.224 | 0.443 | 0.84 | 0.63 | 0.67 | 0.53 | 0.77 | 0.73 |
| 18 | 20621714 | rs6417086 | 2.174 | 1.710 | 2.192 | 2.136 | 0.106 | 0.224 | 0.224 | 0.323 | 0.84 | 0.61 | 0.66 | 0.51 | 0.77 | 0.70 |
| 18 | 23548832 | rs651871 | 2.165 | 0.432 | 1.716 | 0.222 | 0.023 | 0.067 | 0.067 | 0.694 | 0.88 | 0.57 | 0.73 | 0.85 | 0.90 | 0.88 |
| 18 | 26678619 | rs1819942 | 2.440 | 0.490 | 2.135 | 1.955 | 0.012 | 0.049 | 0.049 | 3.006 | 0.72 | 0.41 | 0.64 | 0.56 | 0.67 | 0.60 |
| 18 | 26719357 | rs12605558 | 2.989 | 0.936 | 2.813 | 1.856 | 0.011 | 0.022 | 0.022 | 2.332 | 0.69 | 0.37 | 0.53 | 0.50 | 0.61 | 0.54 |
| 18 | 36946848 | rs1367840 | 2.058 | 0.938 | 0.668 | 1.067 | 0.007 | 0.020 | 0.020 | 0.005 | 0.51 | 0.44 | 0.50 | 0.41 | 0.46 | 0.43 |
| 18 | 71232469 | rs4892113 | 2.017 | 2.933 | 0.162 | 1.115 | 0.001 | 0.003 | 0.003 | 0.173 | 0.59 | 0.57 | 0.61 | 0.50 | 0.45 | 0.45 |
| 19 | 11358440 | rs2116877 | 2.785 | 0.783 | 2.934 | 2.632 | 0.113 | 0.072 | 0.072 | 0.756 | 0.90 | 0.61 | 0.94 | 0.93 | 0.90 | 0.92 |
| 19 | 11536649 | rs34103 | 2.028 | 0.240 | 2.216 | 1.189 | 0.040 | 0.067 | 0.067 | 0.845 | 0.91 | 0.55 | 0.95 | 0.87 | 0.95 | 0.95 |
| 19 | 16591464 | rs9305079 | 2.990 | 2.208 | 2.564 | 0.990 | 0.059 | 0.054 | 0.054 | 0.366 | 0.76 | 0.53 | 0.75 | 0.63 | 0.68 | 0.73 |
| 19 | 16601154 | rs3810196 | 2.781 | 2.253 | 2.683 | 1.014 | 0.050 | 0.066 | 0.066 | 0.351 | 0.75 | 0.46 | 0.61 | 0.60 | 0.67 | 0.72 |
| 19 | 16601194 | rs3810198 | 3.004 | 2.231 | 2.618 | 0.999 | 0.046 | 0.044 | 0.044 | 0.409 | 0.76 | 0.57 | 0.75 | 0.63 | 0.68 | 0.73 |
| 19 | 18964442 | rs2238655 | 2.609 | 1.972 | 0.216 | 1.024 | 0.035 | 0.115 | 0.115 | 0.010 | 0.45 | 0.25 | 0.51 | 0.17 | 0.39 | 0.21 |
| 19 | 29600569 | rs4804862 | 2.412 | 0.118 | 0.553 | 1.733 | 0.007 | 0.051 | 0.051 | 0.102 | 0.38 | 0.11 | 0.24 | 0.35 | 0.38 | 0.37 |
| 19 | 49166875 | rs371690 | 2.398 | 0.104 | 1.776 | 1.415 | 0.003 | 0.013 | 0.013 | 0.661 | 0.72 | 0.34 | 0.46 | 0.56 | 0.64 | 0.68 |
| 20 | 13835998 | rs4814267 | 2.231 | 1.457 | 0.363 | 0.901 | 0.058 | 0.054 | 0.054 | 0.099 | 0.34 | 0.09 | 0.15 | 0.28 | 0.21 | 0.32 |
| 20 | 21375826 | rs6035868 | 3.675 | 0.765 | 2.847 | 2.869 | 0.025 | 0.038 | 0.038 | 0.059 | 0.68 | 0.48 | 0.60 | 0.58 | 0.59 | 0.63 |
| 20 | 48383684 | rs651195 | 2.060 | 0.190 | 1.130 | 1.943 | 0.013 | 0.053 | 0.053 | 0.930 | 0.55 | 0.22 | 0.35 | 0.31 | 0.56 | 0.55 |
| 20 | 49324381 | rs868846 | 2.299 | 2.527 | 2.659 | 0.572 | 0.059 | 0.032 | 0.032 | 0.012 | 0.85 | 0.58 | 0.82 | 0.83 | 0.80 | 0.76 |
| 20 | 49367076 | rs6020721 | 2.504 | 3.446 | 1.967 | 0.075 | 0.103 | 0.067 | 0.067 | 5.264 | 0.52 | 0.13 | 0.46 | 0.66 | 0.34 | 0.51 |
| 20 | 54696043 | rs13037836 | 2.792 | 1.139 | 2.424 | 1.649 | 0.007 | 0.063 | 0.063 | 1.231 | 0.77 | 0.35 | 0.78 | 0.54 | 0.76 | 0.70 |
| 20 | 54699432 | rs989066 | 2.804 | 1.141 | 2.666 | 1.647 | 0.020 | 0.090 | 0.090 | 0.053 | 0.81 | 0.68 | 0.81 | 0.54 | 0.76 | 0.70 |
| 21 | 21168899 | rs467644 | 2.156 | 0.897 | 0.778 | 1.270 | 0.033 | 0.161 | 0.161 | 0.644 | 0.85 | 0.44 | 0.87 | 0.71 | 0.88 | 0.86 |
| 21 | 26605567 | rs8131339 | 2.390 | 1.320 | 2.369 | 1.165 | 0.027 | 0.038 | 0.038 | 0.658 | 0.86 | 0.51 | 0.93 | 0.85 | 0.92 | 0.80 |
| 21 | 30239378 | rs4817258 | 2.509 | 0.600 | 2.018 | 1.033 | 0.008 | 0.175 | 0.175 | 3.095 | 0.55 | 0.12 | 0.45 | 0.18 | 0.59 | 0.55 |
| 21 | 30244170 | rs8134005 | 2.497 | 0.598 | 2.159 | 1.056 | 0.021 | 0.103 | 0.103 | 3.434 | 0.55 | 0.16 | 0.45 | 0.25 | 0.59 | 0.55 |
| 21 | 30527999 | rs2251087 | 2.715 | 1.131 | 0.733 | 0.714 | 0.017 | 0.135 | 0.135 | 1.297 | 0.39 | 0.11 | 0.31 | 0.13 | 0.34 | 0.31 |
| 21 | 30848368 | rs914136 | 2.092 | 0.558 | 1.684 | 1.192 | 0.068 | 0.214 | 0.214 | 1.441 | 0.93 | 0.43 | 0.90 | 0.82 | 0.97 | 0.96 |
| 21 | 30848392 | rs2832341 | 2.120 | 0.556 | 1.683 | 1.192 | 0.068 | 0.208 | 0.208 | 1.907 | 0.93 | 0.44 | 0.91 | 0.84 | 0.97 | 0.96 |
| 21 | 30861858 | rs2211779 | 2.240 | 0.542 | 1.975 | 1.250 | 0.085 | 0.244 | 0.244 | 1.451 | 0.93 | 0.47 | 0.91 | 0.82 | 0.97 | 0.96 |
| 21 | 30865282 | rs980007 | 2.242 | 0.538 | 2.454 | 1.248 | 0.069 | 0.215 | 0.215 | 1.665 | 0.93 | 0.46 | 0.91 | 0.82 | 0.97 | 0.96 |
| 22 | 31917415 | rs4820980 | 2.555 | 0.608 | 0.136 | 1.227 | 0.010 | 0.077 | 0.077 | 1.570 | 0.73 | 0.51 | 0.67 | 0.60 | 0.74 | 0.59 |
| 22 | 35724659 | rs138785 | 2.024 | 0.132 | 0.763 | 2.375 | 0.008 | 0.045 | 0.045 | 0.867 | 0.58 | 0.32 | 0.51 | 0.25 | 0.64 | 0.58 |
| 22 | 37093433 | rs4141439 | 2.678 | 0.135 | 1.306 | 0.485 | 0.034 | 0.032 | 0.032 | 0.041 | 0.69 | 0.18 | 0.62 | 0.79 | 0.76 | 0.72 |
| 22 | 47182944 | rs5769152 | 2.854 | 1.429 | 2.247 | 0.833 | 0.005 | 0.012 | 0.012 | 0.632 | 0.70 | 0.39 | 0.57 | 0.62 | 0.61 | 0.54 |
| 22 | 47198112 | rs710121 | 2.562 | 1.089 | 2.419 | 0.828 | 0.006 | 0.014 | 0.014 | 0.013 | 0.73 | 0.68 | 0.64 | 0.63 | 0.64 | 0.56 |
| 22 | 47200891 | rs6007968 | 2.918 | 1.092 | 2.476 | 0.840 | 0.010 | 0.030 | 0.030 | 0.035 | 0.71 | 0.45 | 0.61 | 0.62 | 0.63 | 0.61 |

Table S3. The 100kb window regions and the number of SNPs under putative selection.

| CHR | MinBP | MaxBP | No. of SNPs | CHR | MinBP | MaxBP | No. of SNPs |
|---|---|---|---|---|---|---|---|
| 7 | 86700000 | 87100000 | 12 | 4 | 32600000 | 32700000 | 1 |
| 11 | 83700000 | 84000000 | 12 | 4 | 108600000 | 108700000 | 1 |
| 8 | 9300000 | 9700000 | 7 | 4 | 142900000 | 143000000 | 1 |
| 7 | 125900000 | 126200000 | 6 | 4 | 152800000 | 152900000 | 1 |
| 11 | 45600000 | 45700000 | 6 | 4 | 153500000 | 153600000 | 1 |
| 11 | 128800000 | 129000000 | 6 | 5 | 3300000 | 3400000 | 1 |
| 17 | 53800000 | 53900000 | 6 | 5 | 11500000 | 11600000 | 1 |
| 16 | 81000000 | 81200000 | 5 | 5 | 20700000 | 20800000 | 1 |
| 2 | 123600000 | 123800000 | 4 | 5 | 24600000 | 24700000 | 1 |
| 3 | 1000000 | 1100000 | 4 | 5 | 54700000 | 54800000 | 1 |
| 3 | 8900000 | 9000000 | 4 | 5 | 57200000 | 57300000 | 1 |
| 4 | 24100000 | 24300000 | 4 | 5 | 108900000 | 109000000 | 1 |
| 8 | 29900000 | 30100000 | 4 | 5 | 125600000 | 125700000 | 1 |
| 14 | 63500000 | 63700000 | 4 | 5 | 145700000 | 145800000 | 1 |
| 15 | 99600000 | 99800000 | 4 | 5 | 165500000 | 165600000 | 1 |
| 17 | 70900000 | 71100000 | 4 | 5 | 165900000 | 166000000 | 1 |
| 21 | 30800000 | 30900000 | 4 | 6 | 32200000 | 32300000 | 1 |
| 1 | 3300000 | 3400000 | 3 | 6 | 35000000 | 35100000 | 1 |
| 2 | 44600000 | 44900000 | 3 | 6 | 40800000 | 40900000 | 1 |
| 2 | 76800000 | 76900000 | 3 | 6 | 81300000 | 81400000 | 1 |
| 2 | 107700000 | 107800000 | 3 | 6 | 84900000 | 85000000 | 1 |
| 3 | 118800000 | 119000000 | 3 | 6 | 167200000 | 167300000 | 1 |
| 4 | 180000000 | 180100000 | 3 | 7 | 1900000 | 2000000 | 1 |
| 7 | 119900000 | 120100000 | 3 | 7 | 8200000 | 8300000 | 1 |
| 8 | 139600000 | 139800000 | 3 | 7 | 8500000 | 8600000 | 1 |
| 8 | 139900000 | 140100000 | 3 | 7 | 12100000 | 12200000 | 1 |
| 11 | 15500000 | 15700000 | 3 | 7 | 14400000 | 14500000 | 1 |
| 11 | 129100000 | 129200000 | 3 | 7 | 16800000 | 16900000 | 1 |
| 12 | 125100000 | 125200000 | 3 | 7 | 19400000 | 19500000 | 1 |
| 13 | 46000000 | 46200000 | 3 | 7 | 21200000 | 21300000 | 1 |
| 13 | 72500000 | 72600000 | 3 | 7 | 73500000 | 73600000 | 1 |
| 13 | 108300000 | 108400000 | 3 | 7 | 78800000 | 78900000 | 1 |
| 15 | 68600000 | 68700000 | 3 | 7 | 98000000 | 98100000 | 1 |
| 16 | 1800000 | 2000000 | 3 | 7 | 135400000 | 135500000 | 1 |
| 19 | 16500000 | 16700000 | 3 | 7 | 137600000 | 137700000 | 1 |
| 22 | 47100000 | 47300000 | 3 | 7 | 147500000 | 147600000 | 1 |
| 1 | 48200000 | 48400000 | 2 | 8 | 17600000 | 17700000 | 1 |
| 1 | 50600000 | 50800000 | 2 | 8 | 18600000 | 18700000 | 1 |
| 1 | 71200000 | 71400000 | 2 | 8 | 24100000 | 24200000 | 1 |
| 1 | 84700000 | 84800000 | 2 | 8 | 33800000 | 33900000 | 1 |
| 1 | 208900000 | 209000000 | 2 | 8 | 108500000 | 108600000 | 1 |
| 2 | 38000000 | 38100000 | 2 | 8 | 116700000 | 116800000 | 1 |
| 2 | 158600000 | 158700000 | 2 | 9 | 900000 | 1000000 | 1 |

Table S3. The 100kb window regions and the number of SNPs under putative selection - Continued

| CHR | MinBP | MaxBP | No. of SNPs | CHR | MinBP | MaxBP | No. of SNPs |
|---|---|---|---|---|---|---|---|
| 2 | 181700000 | 181800000 | 2 | 9 | 14500000 | 14600000 | 1 |
| 2 | 194700000 | 194800000 | 2 | 9 | 27800000 | 27900000 | 1 |
| 3 | 25300000 | 25400000 | 2 | 9 | 32600000 | 32700000 | 1 |
| 3 | 41500000 | 41600000 | 2 | 9 | 32900000 | 33000000 | 1 |
| 3 | 88000000 | 88200000 | 2 | 9 | 79400000 | 79500000 | 1 |
| 4 | 140600000 | 140700000 | 2 | 9 | 87900000 | 88000000 | 1 |
| 4 | 162800000 | 163000000 | 2 | 9 | 114700000 | 114800000 | 1 |
| 5 | 38100000 | 38200000 | 2 | 9 | 123400000 | 123500000 | 1 |
| 5 | 110500000 | 110600000 | 2 | 10 | 44400000 | 44500000 | 1 |
| 5 | 145200000 | 145300000 | 2 | 10 | 80500000 | 80600000 | 1 |
| 6 | 14800000 | 14900000 | 2 | 10 | 90000000 | 90100000 | 1 |
| 6 | 28500000 | 28700000 | 2 | 10 | 108200000 | 108300000 | 1 |
| 6 | 53400000 | 53500000 | 2 | 10 | 119900000 | 120000000 | 1 |
| 6 | 135700000 | 135900000 | 2 | 10 | 120800000 | 120900000 | 1 |
| 7 | 27100000 | 27200000 | 2 | 10 | 121900000 | 122000000 | 1 |
| 7 | 50700000 | 50800000 | 2 | 10 | 128800000 | 128900000 | 1 |
| 7 | 79000000 | 79100000 | 2 | 10 | 131200000 | 131300000 | 1 |
| 7 | 89300000 | 89500000 | 2 | 11 | 20400000 | 20500000 | 1 |
| 9 | 90800000 | 90900000 | 2 | 11 | 20700000 | 20800000 | 1 |
| 10 | 30900000 | 31000000 | 2 | 11 | 22600000 | 22700000 | 1 |
| 10 | 78300000 | 78400000 | 2 | 11 | 34200000 | 34300000 | 1 |
| 10 | 108400000 | 108500000 | 2 | 11 | 59200000 | 59300000 | 1 |
| 10 | 133500000 | 133600000 | 2 | 11 | 67400000 | 67500000 | 1 |
| 11 | 56800000 | 57000000 | 2 | 11 | 76800000 | 76900000 | 1 |
| 11 | 67000000 | 67100000 | 2 | 11 | 84900000 | 85000000 | 1 |
| 11 | 107900000 | 108100000 | 2 | 11 | 88400000 | 88500000 | 1 |
| 12 | 11100000 | 11200000 | 2 | 11 | 102400000 | 102500000 | 1 |
| 12 | 11800000 | 11900000 | 2 | 11 | 104700000 | 104800000 | 1 |
| 12 | 83300000 | 83500000 | 2 | 11 | 130000000 | 130100000 | 1 |
| 12 | 104900000 | 105000000 | 2 | 12 | 4900000 | 5000000 | 1 |
| 12 | 121300000 | 121500000 | 2 | 12 | 76300000 | 76400000 | 1 |
| 14 | 89400000 | 89500000 | 2 | 12 | 81500000 | 81600000 | 1 |
| 15 | 61800000 | 62000000 | 2 | 12 | 109600000 | 109700000 | 1 |
| 15 | 70900000 | 71000000 | 2 | 13 | 21500000 | 21600000 | 1 |
| 15 | 73200000 | 73300000 | 2 | 13 | 22600000 | 22700000 | 1 |
| 16 | 63000000 | 63200000 | 2 | 13 | 67900000 | 68000000 | 1 |
| 17 | 62300000 | 62500000 | 2 | 13 | 72000000 | 72100000 | 1 |
| 18 | 26600000 | 26800000 | 2 | 13 | 83200000 | 83300000 | 1 |
| 20 | 49300000 | 49400000 | 2 | 13 | 93800000 | 93900000 | 1 |
| 20 | 54600000 | 54700000 | 2 | 14 | 85500000 | 85600000 | 1 |
| 21 | 30200000 | 30300000 | 2 | 14 | 100900000 | 101000000 | 1 |
| 1 | 18000000 | 18100000 | 1 | 15 | 72600000 | 72700000 | 1 |
| 1 | 22200000 | 22300000 | 1 | 15 | 95700000 | 95800000 | 1 |
| 1 | 69000000 | 69100000 | 1 | 15 | 97200000 | 97300000 | 1 |

Table S3. The 100kb window regions and the number of SNPs under putative selection - Continued

| CHR | MinBP | MaxBP | No. of SNPs | CHR | MinBP | MaxBP | No. of SNPs |
|---|---|---|---|---|---|---|---|
| 1 | 87200000 | 87300000 | 1 | 16 | 7800000 | 7900000 | 1 |
| 1 | 88900000 | 89000000 | 1 | 16 | 80100000 | 80200000 | 1 |
| 1 | 101500000 | 101600000 | 1 | 17 | 77400000 | 77500000 | 1 |
| 1 | 107700000 | 107800000 | 1 | 18 | 10300000 | 10400000 | 1 |
| 1 | 108400000 | 108500000 | 1 | 18 | 20400000 | 20500000 | 1 |
| 1 | 111200000 | 111300000 | 1 | 18 | 20600000 | 20700000 | 1 |
| 1 | 111500000 | 111600000 | 1 | 18 | 23500000 | 23600000 | 1 |
| 1 | 194200000 | 194300000 | 1 | 18 | 36900000 | 37000000 | 1 |
| 1 | 203400000 | 203500000 | 1 | 18 | 71200000 | 71300000 | 1 |
| 1 | 231500000 | 231600000 | 1 | 19 | 11300000 | 11400000 | 1 |
| 1 | 231800000 | 231900000 | 1 | 19 | 11500000 | 11600000 | 1 |
| 1 | 248100000 | 248200000 | 1 | 19 | 18900000 | 19000000 | 1 |
| 2 | 21800000 | 21900000 | 1 | 19 | 29600000 | 29700000 | 1 |
| 2 | 158200000 | 158300000 | 1 | 19 | 49100000 | 49200000 | 1 |
| 2 | 192600000 | 192700000 | 1 | 20 | 13800000 | 13900000 | 1 |
| 2 | 218200000 | 218300000 | 1 | 20 | 21300000 | 21400000 | 1 |
| 2 | 234200000 | 234300000 | 1 | 20 | 48300000 | 48400000 | 1 |
| 3 | 10600000 | 10700000 | 1 | 21 | 21100000 | 21200000 | 1 |
| 3 | 29500000 | 29600000 | 1 | 21 | 26600000 | 26700000 | 1 |
| 3 | 33100000 | 33200000 | 1 | 21 | 30500000 | 30600000 | 1 |
| 3 | 109700000 | 109800000 | 1 | 22 | 31900000 | 32000000 | 1 |
| 3 | 137500000 | 137600000 | 1 | 22 | 35700000 | 35800000 | 1 |
| 4 | 16500000 | 16600000 | 1 | 22 | 37000000 | 37100000 | 1 |

Table S4. The Gene Ontology analysis* results after Bonferroni corrections.

| GO biological process complete | Number (Background) | Number (Input) | Expected | Fold Enrichment | +/- | p-value |
|---|---|---|---|---|---|---|
| Glycosaminoglycan biosynthetic process | 100 | 11 | 1.39 | 7.91 | + | 0.003 |
| **a.** Aminoglycan biosynthetic process | 105 | 11 | 1.46 | 7.53 | + | 0.005 |
| **b.** Aminoglycan metabolic process | 161 | 12 | 2.24 | 5.36 | + | 0.044 |
| **c.** Glycosaminoglycan metabolic process | 151 | 12 | 2.10 | 5.71 | + | 0.024 |

***The list of genes used as input for GO analysis**

*ABCB4, ABTB2, ACACB, ACAT1, ACSS3, ACTL8, ACVR1, ACY3, ADAM28, AGR2, AGR3, AHI1, ALDH3B2, ANGPT1, ANGPTL8, ANKRD13D, ANKS1A, APLP2, APTX, ARHGAP32, ARHGEF10L, ARHGEF16, ATG16L1, ATM, ATMIN, ATP2B2, ATP2B2-IT2, B4GALT4, B4GALT4-AS1, B4GALT5, BACH1-IT3, BAIAP2L1, C12orf43, C16orf46, C19orf44, C3orf30, C3orf38, C6orf10, CA11, CACNG2, CALR3, CAMK4, CAMKMT, CAPN5, CASC2, CASP12, CAVIN2, CCDC151, CDH10, CELF6, CENPN, CEP162, CERK, CERS1, CGGBP1, CHERP, CHST1, CHST11, CMC2, CNTNAP2, COG3, COL22A1, COMP, CREB3L2, CROT, CRTAP, CTC-510F12.2, CTNND2, CUL5, CYTIP, DACH1, DBP, DCTN6, DDX5, DEPDC1, DGKB, DGKD, DIRC3, DIRC3-AS1, DISC1, DLG2, DMRT1, DMRT3, DMTF1, DOCK1, DOCK6, EGLN1, EIF3A, EIF4H, ELAVL3, ELAVL4, EME2, EPS15L1, ERICH6B, ETV6, FAHD1, FAM133DP, FAM155A, FAM180A, FAM45A, FAM83E, FANCF, FSTL5, FUT2, GAS2, GCLC, GCSH, GDF1, GLB1, GPC6, GPX5, GRAMD2B, GRAMD3, GRB10, GRK2, GRM5, GRM8, GRXCR2, HAGH, HAS1, HEXA, HEXA-AS1, HMOX1, HNF1A, HOTAIRM1, HOXA-AS2, HOXA-AS3, HOXA1, HOXA2, HOXA3, HOXA4, HOXA5, HOXA6, HOXA7, HS3ST6, HSP90B2P, HSPG2, HTATIP2, HTR1F, ICA1, IGFALS, IGSF11, INPP4B, ITGA11, KANK2, KCNA3, KCNA6, KCND2, KCNH5, KDM2A, KIAA1009, LATS2, LDB2, LEPROTL1, LIMK1, LINC00189, LINC00211, LINC00254, LINC00271, LINC00364, LINC00367, LINC00533, LINC00841, LINC00913, LINC01210, LINC01564, LINC01717, LINC01788, LINC02107, LINC02119, LINC02124, LINC02241, LOC100130880, LOC100130987, LOC100505685, LOC100505938, LOC100507274, LOC101927136, LOC101927154, LOC101927171, LOC101927244, LOC101927420, LOC101927488, LOC101927633, LOC101927760, LOC101927879, LOC101928052, LOC101928186, LOC101928357, LOC101928396, LOC101928828, LOC101928850, LOC101928888, LOC101928901, LOC101929163, LOC101929230, LOC101929294, LOC101929501, LOC101929555, LOC101929559, LOC101929727, LOC101929745, LOC102606465, LOC102724084, LOC102724957, LOC105370888, LOC105373021, LOC399975, LOC643733, LRIF1, LRMDA, LRRC28, LRRC55, LTN1, LYZL2, MAD1L1, MAGI2, MAGI2-AS3, MAML3, MAP3K7CL, MAPK8IP3, MBOAT4, MCM5, MED26, MEGF9, MEIOB, MGMT, MGST2, MILR1, MIR3064, MIR3909, MIR3922, MIR4699, MIR5047, MIR597, MIR6069, MIR7852, MIR924HG, MMP20, MMP7, MRPS34, MSRB1, MTREX, MTUS1, MYO7A, N6AMT1, NELL1, NKX2-4, NME3, NPAT, NTN5, NTNG1, NUBP2, NXPH1, OASL, OMP, OPTC, OR2AK2, OR2L13, OR2L1P, OR2L2, OR2L5, OR2L8, OR4D10, OR4D11, OR4D6, OR4D9, OR5A1, OR5AK4P, PAPSS1, PARD6B, PCA3, PCTP, PECAM1, PKD1L2, PLPP1, POLG2, POU4F3, PPARGC1A, PRDM16, PRELID2, PRELP, PRH1, PRH1-PRR4, PRH1-TAS2R14, PRKACB, PRKCSH, PRMT3, PRUNE2, PSD3, PTGER3, RAD18, RARB, RBBP8, RBFOX3, RBMS3, RGL3, RHOJ, RIPOR3, RNLS, RP1-27K12.2, RP11-497K21.1, RP11-624C23.1, RP11-76N22.2, RP4-694A7.4, RP5-952N6.1, RPL18, RPL3L, RPS6KA2, SAG, SALRNA2, SALRNA3, SAMD13, SARAF, SCHLAP1, SDPR, SEC1P, SEL1L2, SFI1, SFXN4, SH3GLB1, SKIV2L2, SLC13A4, SLC30A7, SLC35E1, SLC39A11, SNORA19, SORCS1, SPACA4, SPHK2, SPPL3, SPRTN, SPSB3, SS18, SSH3, ST14, SULT2B1, SUSD5, SVILP1, SYNM, TAF1L, TAS2R19, TAS2R20, TAS2R31, TAS2R50, TBC1D22A, TBX10, TCP11, TEX2, TMEM100, TMEM154, TMEM202, TMEM202-AS1, TMEM243, TMEM66, TMPPE, TMTC2, TNKS, TOM1, TP53AIP1, TP53TG1, TRABD2B, TRPS1, TSNAX-DISC1, TSPAN13, TTC23, UACA, ULK4, UPF1, UPK1B, UQCRFS1, VAV3, WDR25, XLOC_009911, XRN2, ZBED9, ZBTB44, ZNF653, ZNF654.*

Table S5. Regions under selection in the Kuwait population and associated disease phenotypes.

| CHR | Region (bp) | Number of SNPs | Genes* | Diseases and Disorders** | PheWAS Traits*** |
|---|---|---|---|---|---|
| 7 | 86700000-87100000 | 12 | *BC035377, TMEM243, CROT, ABCB4* | Cholelithiasis (Rosmorduc et al. 2003); Cholestatis (Lang et al. 2007) | N/A |
| 11 | 83700000-84000000 | 12 | *DLG2* | Body Height (Estrada et al. 2009); Calcium-Binding Proteins (Benjamin et al. 2007); Chemokine CCL2; Protein quantitative trait loci (Melzer et al. 2008); Cystatins (Hwang et al. 2007); Echocardiography (Vasan et al. 2007); Hip (Kiel et al. 2007); Parkinson's Disease (Fung et al. 2006); Phospholipids (Demirkan et al. 2012) | N/A |
| 8 | 9300000-9700000 | 7 | *TNKS* | Adiposity (Lindgren et al. 2009); Asthma (Ober et al. 2000) | hypertension; Platelet distribution width; Standing height; Body mass index (BMI); Red blood cell (erythrocyte) distribution width; Arm fat percentage (left); Arm fat percentage (right); Impedance of leg (left); Impedance of leg (right); Arm fat mass (left); Arm fat mass (right); Cheese intake; Eosinophill percentage; Impedance of arm (right); Impedance of whole body; Leg fat mass (left); Leg fat mass (right) |
| 7 | 125900000-126200000 | 6 | *GRM8* | Carotid Artery Diseases; Carotid atherosclerosis in HIV infection (Shrestha et al. 2010); Chemokines (Bozaoglu et al. 2010); Depression (Terracciano et al. 2010); Electrocardiography (Newton-Cheh et al. 2007); schizophrenia (Takaki et al. 2004); Triglycerides (Kathiresan et al. 2007) | N/A |
| 11 | 45600000-45700000 | 6 | *CHST1* | N/A | N/A |
| 11 | 128800000-129000000 | 6 | *TP53AIP1, ARHGAP32* | Cholesterol, LDL (Kathiresan et al. 2007) | Mean platelet (thrombocyte) volume |
| 17 | 53800000-53900000 | 6 | *PCTP* | Marijuana Abuse (Agrawal et al. 2011) | N/A |
| 16 | 81000000-81200000 | 5 | *ATMIN, C16orf46, GCSH* | Attention deficit hyperactivity disorder and conduct disorder (Anney et al. 2008); Body Weights and Measures (Fox et al. 2007); smoking cessation (Uhl et al. 2010) | Mean reticulocyte volume; Mean corpuscular volume; Mean sphered cell volume |

* Harboring at least one of the selected SNPs, according to UCSC Genome Browser;

** According to UCSC Genome Browser;

*** Threshold: -log10 (p-value) > 8, according to GeneAtlas.

Table S6. The PheWAS traits of the seven SNPs within the putatively selected haplotype.

| PheWAS Trait | rs6987057 | rs9644677 | rs9329203 | rs13273033 | rs4841196 | rs13276086 | rs12545912 | No. of SNPs |
|---|---|---|---|---|---|---|---|---|
| Hypertension | * | 9.4804 | 9.4727 | 10.0742 | 11.8367 | 14.2411 | 10.1253 | 6 |
| Platelet distribution width | 23.7242 | * | 9.0920 | 8.3402 | 12.6545 | 29.6079 | 8.6918 | 6 |
| Standing height | * | * | 8.9109 | 10.2032 | * | 15.8442 | 10.6244 | 4 |
| Body mass index (BMI) | 8.1483 | * | * | * | 10.9372 | 17.4981 | * | 3 |
| Red blood cell (erythrocyte) distribution width | 9.7441 | * | * | * | 9.2074 | 14.3671 | * | 3 |
| Arm fat percentage (left) | * | * | * | * | 9.0920 | 13.1893 | * | 2 |
| Arm fat percentage (right) | * | * | * | * | 8.6213 | 12.6453 | * | 2 |
| Impedance of leg (left) | * | * | * | * | 10.3407 | 19.2792 | * | 2 |
| Impedance of leg (right) | * | * | * | * | 8.4599 | 16.8135 | * | 2 |
| Arm fat mass (left) | * | * | * | * | * | 10.6233 | * | 1 |
| Arm fat mass (right) | * | * | * | * | * | 10.6568 | * | 1 |
| Cheese intake | * | * | * | * | * | 8.0460 | * | 1 |
| Eosinophill percentage | * | * | * | * | * | 8.8225 | * | 1 |
| Impedance of arm (right) | * | * | * | * | * | 8.2298 | * | 1 |
| Impedance of whole body | * | * | * | * | * | 13.6763 | * | 1 |
| Leg fat mass (left) | * | * | * | * | * | 9.2310 | * | 1 |
| Leg fat mass (right) | * | * | * | * | * | 9.6228 | * | 1 |

* = -log10(p-value) < 8 (Threshhold)
